# Probiotic consortia improve anti-viral immunity to SARS-CoV-2 in Ferrets

**DOI:** 10.1101/2021.07.23.453521

**Authors:** J Lehtinen Markus, Kumar Ritesh, Zabel Bryan, M Mäkelä Sanna, Nedveck Derek, Tang Peipei, Latvala Sinikka, Guery Sebastien, R Budinoff Charles

## Abstract

Probiotics have been suggested as one solution to counter detrimental health effects by SARS-CoV-2, however, data so far is scarce. We tested the effect of two probiotic consortia, OL-1 and OL-2, against SARS-CoV-2 in ferrets and assessed their effect on cytokine production and transcriptome in a human monocyte-derived macrophage (Mf) and dendritic cell (DC) model.

The results showed that the consortia significantly reduced the viral load, modulated immune response, and regulated viral receptor expression in ferrets compared to placebo. In human Mf and DC model, OL-1 and OL-2 induced cytokine production and genes related to SARS-CoV-2 anti-viral immunity.

The study results indicate that probiotic stimulation of the ferret immune system leads to improved anti-viral immunity against SARS-COV-2 and that critical genes and cytokines for anti-SARS-CoV-2 immunity are stimulated in human immune cells *in vitro*. The effect of the consortia against SARS-CoV-2 warrants further investigations in human clinical trials.

## INTRODUCTION

The emergence and fast spread of severe acute respiratory syndrome coronavirus 2 (SARS-CoV-2) that causes coronavirus disease 2019 (COVID-19), has resulted in an acute need for effective therapies and vaccination, but also a need for effective dietary strategies for improving immune function. The potential of probiotics, live microorganisms that confer health benefit on the host, in managing SARS-CoV-2 infection has not been extensively studied, despite the well-established connection between gut microorganisms and immune function (Hill et al., 2014).

In humans, COVID-19 symptoms vary from mild respiratory symptoms such as fever, fatigue, and dry cough, to severe respiratory failure with acute respiratory distress symptom, acute cardiac injury and multiorgan failure. Although most individuals subsequently resolve the infection, the disease may also progress to severe pneumonia in susceptible groups. The incubation time of the virus in human is about 5 days, severe disease typically develops 8 days after symptom onset, and death occurs at about 16 days (Schultze and Aschenbrenner, 2021).

In February 2020, the World Health Organization (WHO) assembled an international panel to develop animal models for COVID-19 to accelerate the testing of anti-SARS-CoV-2 therapies (Munoz-Fontela et al., 2020). Ferret model is one of the recommended by the panel to study SARS-CoV-2. Ferrets (*Mustela putorius furo*) have been used traditionally for testing the pathogenicity and transmission of human respiratory viruses, including influenza. Ferrets are highly susceptible to SARS-CoV-2 and show an increase in viral load already one day post infection, transmit the virus efficiently, and shed it in nasal washes, saliva, urine, and feces for 8 days post infection (Kim et al., 2020). Ferrets recapitulate the human-to-human transmission but not the severity of the COVID-19 in humans (Kim et al., 2020).

SARS-CoV-2 uses angiotensin-converting enzyme 2 (ACE2) as the main receptor for cell entry (Hoffmann et al., 2020), but with several co-factors involved, including neuropilin-1 (Cantuti-Castelvetri et al., 2020). The domain in ferret ACE2 that binds the spike protein of SARS-CoV-2 differs by two amino acids from its human homologue and binds to SARS-CoV-2 with similar affinity (Munshi et al., 2021). SARS-CoV-2 primarily infects epithelial cells in the respiratory tract (Alfi et al., 2021), however, infections of the gastrointestinal tract have also been observed (Stanifer et al., 2020), but not extensively studied. In innate immune cells, such as monocytes and macrophages, the infection seems to be abortive (Zheng et al., 2021).

SARS-CoV-2 is an enveloped virus with an +ssRNA genome. The specific innate immune receptors and signaling pathways triggering the interferon (IFN) response in the case of SARS-CoV-2 are yet to be fully determined, however, based on current findings Toll-like receptor (TLR) 3, TLR7, TLR8, RIG-I, and MDA5 recognize viral RNA and drive MyD88 and NF-κB -dependent pro-inflammatory and type I IFN response that further trigger STAT1, STAT2, and IRF9 -dependent induction of IFN-stimulated genes (ISG) (Schultze and Aschenbrenner, 2021). On the other hand, SARS-CoV-2 suppresses the innate immune activation and IFN response to enable its replication in the host cells (Lee et al., 2020; Taefehshokr et al., 2020) that seems to lead to inefficient B cell and T cell responses (Zhou et al., 2020a). Human studies indicate that delayed innate IFN response results at a higher risk to develop severe COVID-19 (Hadjadj et al., 2020). In COVID-19 patients with moderate to severe disease, high chemokine (e.g. CXCL2, CXCL8, CXCL9, CCL2) and pro-inflammatory cytokine (e.g. interleukin (IL)-1β, IL-6, IL-1Ra) response was detected in bronchoalveolar lavage fluid (BALF) (Zhou et al., 2020b), in post-mortem lung biopsies and serum samples (Blanco-Melo et al., 2020) but without significant upregulation of type I or type III IFNs, although ISGs from e.g. IFIT and IFITM families were induced. Accordingly, intranasal IFN-I administered pre-or post-virus challenge reduced disease burden (Hoagland et al., 2021). The above suggest that stimulation of innate immune function could be beneficial in controlling the viral replication and eradication.

In the past two decades the effect of probiotics on immune function and respiratory viral infections have been studied in numerous clinical studies, but with variable quality. Although meta-analyses indicate efficacy of probiotics against acute upper respiratory tract (URT) infections in general (Hao et al., 2015; King et al., 2014; Shi et al., 2021), the results per strain vary, and accordingly probiotic effects on immunity should be investigated on a per strain or consortia basis (Hill et al., 2014). Clinical studies on specific strains like *Bifidobacterium* (B.) *lactis* Bl-04 showed that specific probiotics can reduce the risk of URT illness episodes in healthy adults (West et al., 2014) or reduce rhinovirus load in human nasal wash samples (Turner et al., 2017). The clinical and pre-clinical evidence suggest that use of probiotics could prime immune system prior to viral infection (Lehtoranta et al., 2020). For example, *Lactobacillus* (L.) *acidophilus* NCFM induced the upregulation of TLR3, IL-12, and IFN-β in murine DCs (Weiss et al., 2010) and *Lacticaseibacillus* strains inhibited influenza A replication in human monocyte-derived macrophages (Miettinen et al., 2012).

One of the hypotheses for the probiotic mode of actions against SARS-CoV-2 respiratory and intestinal infection is via strengthening the gut epithelial barrier and beneficial modulation of the gut microbiota and the immune system (Harper et al., 2021). Age and comorbidities that make people more susceptible to severe COVID-19 are associated with perturbed gut microbiota, dysbiosis, and decrease in epithelial barrier integrity in the gut (Vignesh et al., 2021). SARS-CoV-2 infects gut epithelial cells (Stanifer et al., 2020) which may cause lipopolysaccharide (LPS) and other pathogen-associated molecular patterns (PAMPs) to leak across the epithelial barrier. The resulting innate immune activation via the gut-lung-axis may contribute to the cytokine storm in these individuals (Vignesh et al., 2021).

To investigate the effect and mechanism of action of probiotics against SARS-CoV-2 we designed two probiotic consortia to stimulate innate immune function against SARS-CoV-2: OL-1 (B. *lactis* Bl-04, B. *longum* subsp. *infantis* Bi-26, *Lacticaseibacillus rhamnosus* Lr-32, *Lacticaseibacillus paracasei* Lpc-37, *Ligilactobacillus salivarius* Ls-33) and OL-2 (B. *lactis* Bi-07, *L. acidophilus* NCFM, *Limosilactibacillus fermentum* SBS-1, *Lactococcus lactis* Ll-23, *Streptococcus thermophilus* St-21). We tested the effect of the probiotic consortia in a ferret SARS-CoV-2 challenge model and in human monocyte-derived macrophage (Mf) and dendritic cell (DC) model. The results in ferrets show that OL-1 and OL-2 modulate the duodenal and lung immune response during infection, ACE2 expression in duodenum, and reduce SARS-CoV-2 viral load in nasal washes. We further show that the exposure of OL-1 and OL-2 to human monocyte-derived Mfs and DCs results in upregulation of genes and cytokines critical for early innate immune activation against SARS-CoV-2.

## RESULTS

### Probiotic consortia decrease the nasal wash SARS-CoV-2 titers in ferrets

Ferrets have been shown to model SARS-CoV-2 replication and infection in the airways and to express ACE2 (Kim et al., 2020). We evaluated the effect of the two probiotic consortia OL-1 and OL-2 on the SARS-CoV-2 infection, ACE2 expression and immune function in ferrets in two placebo-controlled studies: main (**Figure 1**) and pilot (**Figure S1**).

**Figure 1.**
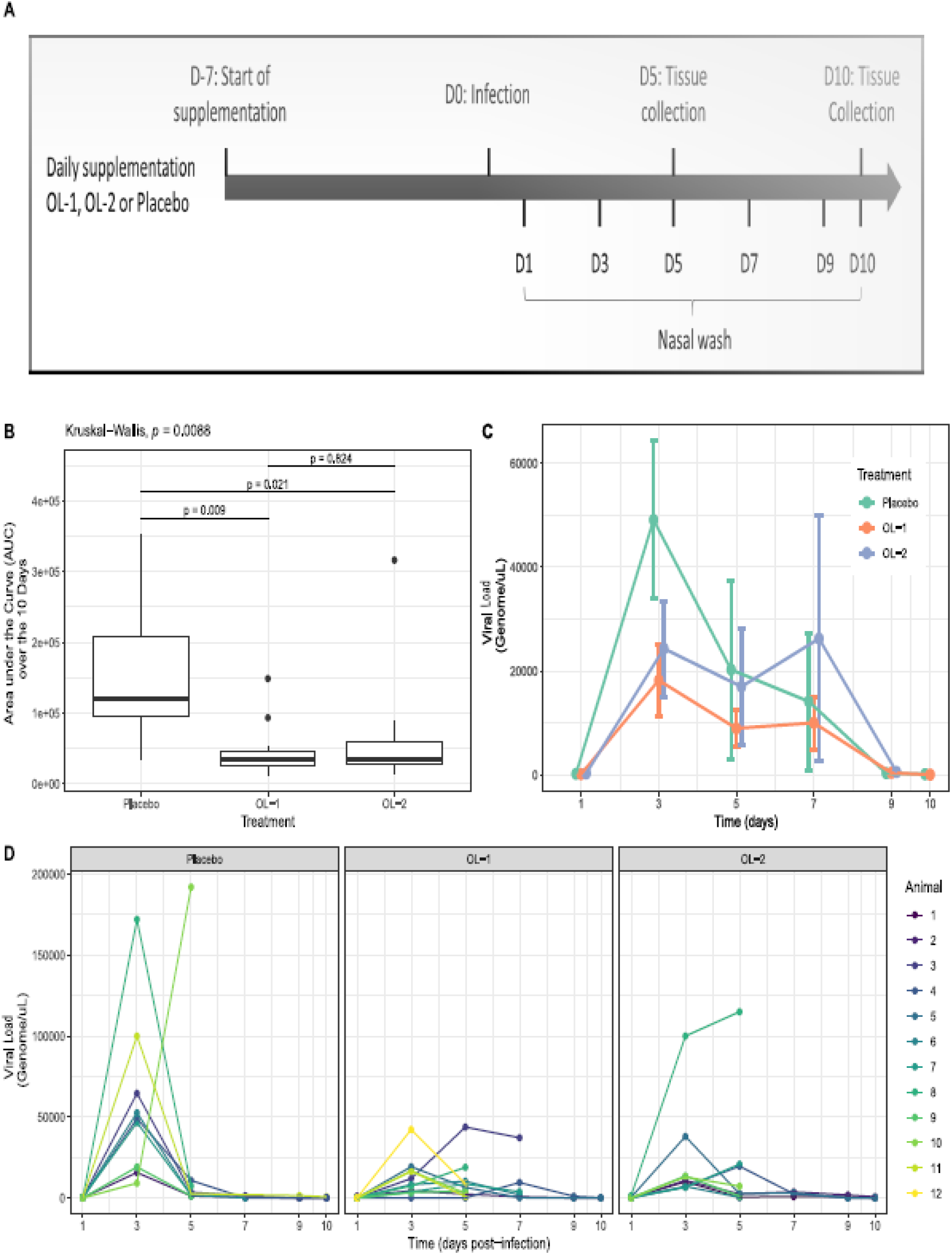
The main ferret study and nasal wash viral load analysis. The ferrets were supplemented with OL-1 or OL-2 or placebo and infected with SARS-CoV-2. (A) The ferret study design and time points for treatment supplementation, SARS-CoV-2 infection, tissue and nasal wash collection (D=Day); (B) The nasal wash viral load AUC analysis (Line: median Box: lower and upper quartile, Whiskers: min and max; Kruskal-Wallis test and pairwise Wilcoxon tests) (C) The nasal wash viral load timepoint analysis (Symbol: mean; error bars: standard error); (D) Time course of individual viral loads in nasal washes from ferrets.

In the main study, probiotic consortia were supplemented 7 days (D) prior (D-7) to viral infection (D0) and until D10 post infection. Lung and duodenum samples were gathered for qPCR gene expression analysis (D5 and D10) and tissue staining (D5 and D10); and nasal washes were collected D1-D10 for determining the viral load by qPCR (**Figure 1A**). The SARS-CoV-2 viral load over the infection time (D1-D10) showed statistically significant differences in the area under the curve (AUC) between the OL-1, OL-2, and placebo groups (p=0.0088; Kruskall-Wallis) (**Figure 1B**). Post-hoc pairwise Wilcoxon tests further demonstrated the significant differences in AUC between the placebo and the OL-1 (p=0.009) and OL-2 (p=0.021) groups, respectively. On average, OL-1 and OL-2 reduced the AUC by 68% and 52%, respectively, as compared to the placebo group, suggesting influence of the probiotic consortia on the anti-viral immunity against SARS-CoV-2.

The time-course analysis of the SARS-CoV-2 nasal wash qPCR showed that the viral loads peaked in general at D3 (average viral titer of 4.9×10^4^ genome/µL, 1.8×10^4^ genome/µL, and 2.4×10^4^ genome/µL in the Placebo, OL-1, and OL-2 groups, respectively) and were mostly resolved by D9 (average viral titer of 319 genome/µL, 394 genome/µL, and 552 genome/µL in the placebo, OL-1, and OL-2 groups, respectively) (**Figure 1C**). Further investigation of the individual line plots (**Figure 1D**) showed that the viral loads of few animals show later peaking (1/9, 5/11, 3/9 animals in the Placebo, OL-1, and OL-2 groups, respectively). Most later peaking animals were found in the treatment groups that could reflect the effects observed on the viral replication.

In comparison to the main study, the pilot study had a different probiotic supplementation regime, where OL-1 was administered 21 days prior to infection (D-21) and OL-2 one day post infection (D1) (**Figure S1**). The viral load analysis showed undetectable to very low viral genomes in the samples (**Figure S1**), despite the use of same viral dose to induce the infection and housing dependent infectivity, indicating potential existing immunity to the virus or issues with the viral inoculum. The pilot study did not show treatment effect on the viral load. However, the lung and duodenum samples at D0 (pre-infection), D5, and D21 were collected for later qPCR analyses, and D0 samples were used later as a comparator for the main study responses.

The body temperature of the ferrets and visual symptoms of viral infection were monitored in both studies. The SARS-CoV-2 infection in ferrets did not affect the body temperature, body weight or induce coughing, sneezing or dyspnea in the animals during infection, in line with previous literature (Peacock et al., 2021; Ryan et al., 2021).

#### Lung and duodenal gene expression show changes by probiotic consortia

To understand further the mechanism of action of OL-1 and OL-2 on the lung viral load decrease, we investigated the effect of the probiotic consortia on the ferret immune system pre-and post-infection. Lung and duodenal tissues from the main (**Figure 1A**) and the pilot study (**Figure S1**) were collected and gene expression measured using qPCR (**Figure 2**). Importantly, pre-infection samples (D0) were available only for the OL-1 and placebo groups from the pilot study. Thus, D0 placebo group samples from the pilot study were used as a baseline to normalize not only the pilot (**Figure 2A**), but also the main study qPCR response data (**Figure 2B**). Statistical analysis indicated only minor changes in the pilot study for lung and duodenal gene expression (**Figure 3A**), in line with the poor infectivity in the experiment. Instead, in the main study, IFN (*IFNA and IFNL1*), cytokine (IL-10) and chemokine (*CCL2, IL8*) gene expression was increased in responses to the SARS-CoV-2 infection in all the treatment groups (adjusted p<0.1; FDR) (**Figure 2B**). It is noteworthy that IFNG expression was downregulated in the lung tissue and ACE2 expression was downregulated in the duodenum, whereas it was upregulated in the lungs (**Figure 2B**). At D5 both consortia, but not placebo, had a significant increase in *CXCL10* in duodenum and lungs (adjusted p<0.1; FDR) (**Figure 2B**).

**Figure 2.**
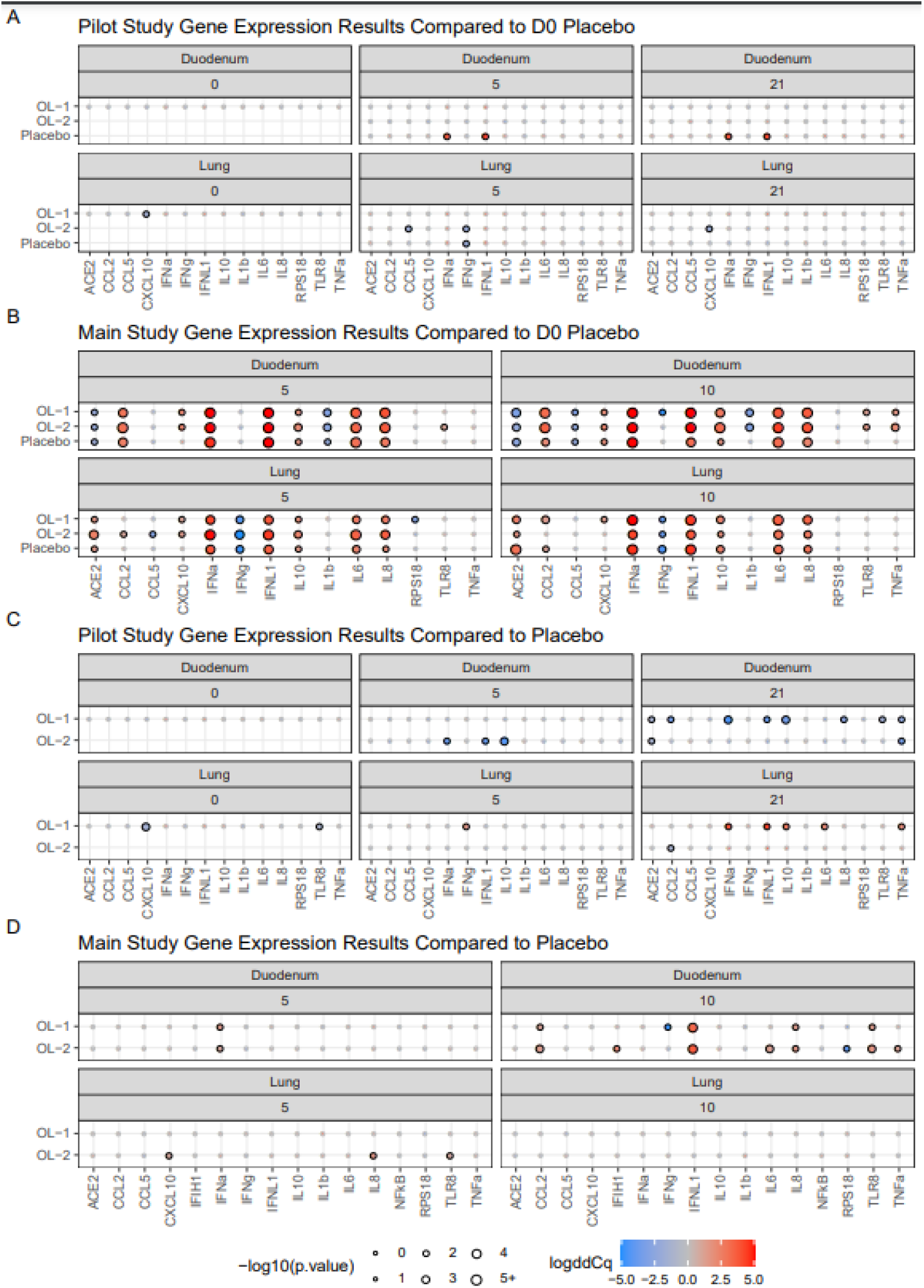
Immune marker gene expression of the ferrets to SARS-CoV-2 and the effect of the treatments. Balloon plots showing gene expression levels as log2 fold changes for the pilot and main studies. Pilot study placebo tissues at D0 prior to viral challenge were used as a comparator for post-infection timepoints in (A) the pilot and (B) the main study. Placebo treatments at the same timepoints for (C) the pilot and (D) the main study was used as the comparator. Color of balloons denotes direction and magnitude of log2 fold change, with blue showing a decreased log2 fold change relative to the placebo, and red and increased log2 fold change. Size of the balloons corresponds to the –log10 of the p-value (adjusted with the Tukey method for a family of estimates) from a post-hoc pairwise comparison of the estimated marginal means to the placebo reference, larger balloons mean a smaller p-value. Statistically significant fold changes are marked by a black outline around the balloon (p-value < 0.1).

**Figure 3.**
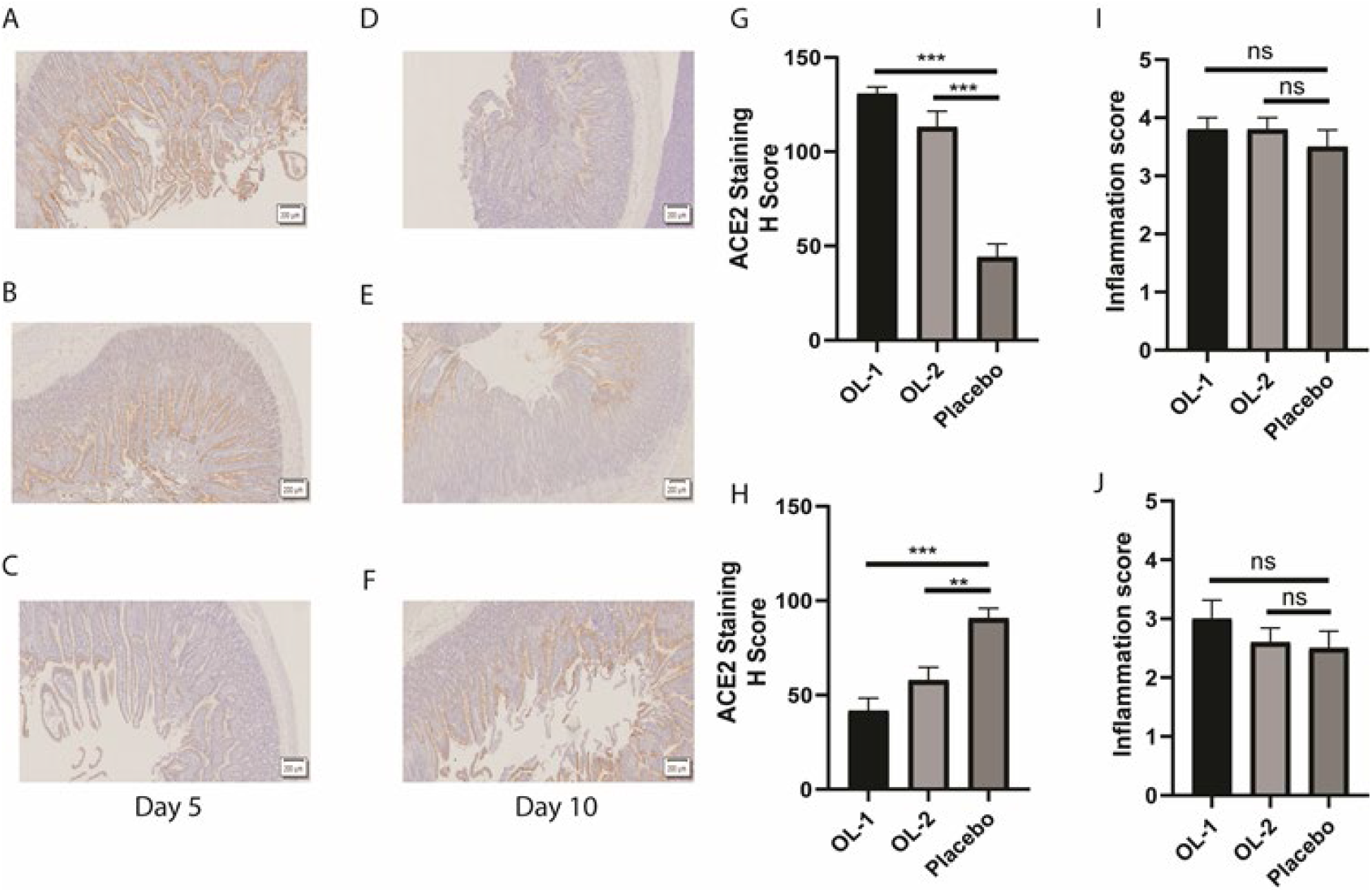
Localization and expression of ACE2 and inflammatory scoring in duodenum. Representative images and quantification of ACE2 immunohistochemical staining from the main study. Panels (A)-(C) are from D5 and (D)-(F) are from D10. (A) and (D) are from OL-1, (B) and (E) are from OL-2 and (C) and (F) are from placebo groups. (G) Quantification of ACE2 expression for D5. (H) Quantification of ACE2 expression for D10. (I) and (J) H&E-stained duodenum sections were evaluated for inflammation (0 no inflammation to 4 severe inflammation) and average inflammation score for each treatment group is shown in (I) for D5 and (J) for D10. Data is presented as mean ± SEM. Statistical analysis was performed using one-way, ANOVA followed by Dunnett’s multiple comparisons test. ns, not significant, **, p < 0.01; ***, p < 0.001.

To compare the effect of OL-1 and OL-2 to placebo, we did a timepoint comparison of the data for the pilot and the main study. In the pilot study there were minor differences at D0 and D5. At pre-infection (D0), OL-1 decreases *CXCL10* and *TLR8* expression in the lung samples (adjusted p<0.1; FDR) (**Figure 2C**), indicating potential influence of the supplementation alone on the lungs. At D5 OL-2 decreased *IFNA, IFNL1*, and *IL10* in duodenum, and OL-1 increased *IFNG* expression in lungs (adjusted p<0.1; FDR) (**Figure 2C**), suggesting influence on the IFN responses. At D21, when the infection was cleared, OL-1 had broadly decreased expression of cytokines and IFNs in the duodenum and an increase in *IFNA, IFNL1, IL10, IL6*, and *TNFA* in lungs (adjusted p<0.1; FDR). Further, both consortia decreased ACE2 expression in the duodenum (adjusted p<0.1; FDR) (**Figure 2C**). The results at D21 could be an indication of post-infection influence of the probiotic consortia on the recovery from the infection.

In the main study, expression of several immune genes was regulated by OL-1 and OL-2 (**Figure 2D**). At D5 both consortia induced upregulation of *IFNA* in the duodenum, and OL-1 further upregulated *CXCL10, IL8*, and *TLR8* in the lungs compared to placebo (adjusted p<0.1; FDR). At D10 both consortia had higher *CCL2, IFNL1, IL8*, and *TLR8* gene expression in duodenum compared to placebo (adjusted p<0.1; FDR). OL-1 had in addition a decrease in *IFNG* expression and OL-2, on the other hand, increased *IFIH1, IL6*, and *TNFA* (adjusted p<0.1; FDR) (**Figure 2D**). In the lungs, there were no significant differences at D10 between the groups. Overall, the results indicate stronger immunomodulation in the duodenum, but also effects in the lungs, in line with the route of administration by gavage. The expression patterns in the duodenum and lungs were quite similar suggesting activation of gut-lung axis by both consortia (**Figure 2B**).

### Probiotic consortia modulate duodenal ACE2 surface expression

We found in the qPCR analysis of the pilot study a decrease of *ACE2* expression at D21 in duodenum (**Figures 2C**). As humans have gastrointestinal symptoms and potential viral replication in the gut, we wanted to further determine by tissue staining if the probiotic supplementation would influence viral receptor ACE2 expression and inflammation in the duodenal tissues (**Figure 3**). For ACE2, during acute infection at D5, OL-1 and OL-2 group tissue sections had increased staining compared to placebo group (p<0.01; ANOVA, Dunnett’s) (**Figure 3A-C and G**), suggesting cytoprotective effects during inflammation. At D10, during the resolution phase of the infection, the ACE2 staining was decreased in OL-1 and OL-2 groups compared to D5 and to placebo (p<0.01; ANOVA, Dunnett’s) (**Figure 3D-F and H**). The results suggest differential effects of the OL-1 and OL-2 compared to placebo on ACE2 cell surface expression over the course of infection. It is noteworthy that as we did not have the pre-infection D0 staining samples for ACE2, the normal level of ACE2 is not known.

The inflammatory scorings of the tissue sections showed severe inflammation in all the treatment groups and gradual resolution by D10 (Average score of 3.5 at D5, and 2.5 at D10 of maximum 4.0 in all groups), supporting the result of the increase in inflammatory gene expression levels by qPCR (**Figure 2B**). However, the fecal pellets remained of normal consistency in the ferrets. The treatments with OL-1 or OL-2 did not have an effect on the duodenal inflammation scores (p>0.05; ANOVA, Dunnett’s).

#### Human macrophages and dendritic cells are stimulated by OL-1 and OL-2

To test how OL-1 and OL-2 could potentially influence anti-SARS-CoV-2 immunity in humans, we used fresh human peripheral blood monocyte-derived macrophages (Mf) and dendritic cells (DC). The immune cells were incubated with OL-1, OL-2, controls, or TLR agonist mix pIC (TLR3) and R848 (TLR7/8) – that mimics immune response to viral RNA. We also used the combination of Poly I:C (pIC)+R848 with OL-1 or OL-2 to evaluate modulation of the cell response to pIC+R848 by the consortia (**Figure 4**).

**Figure 4.**
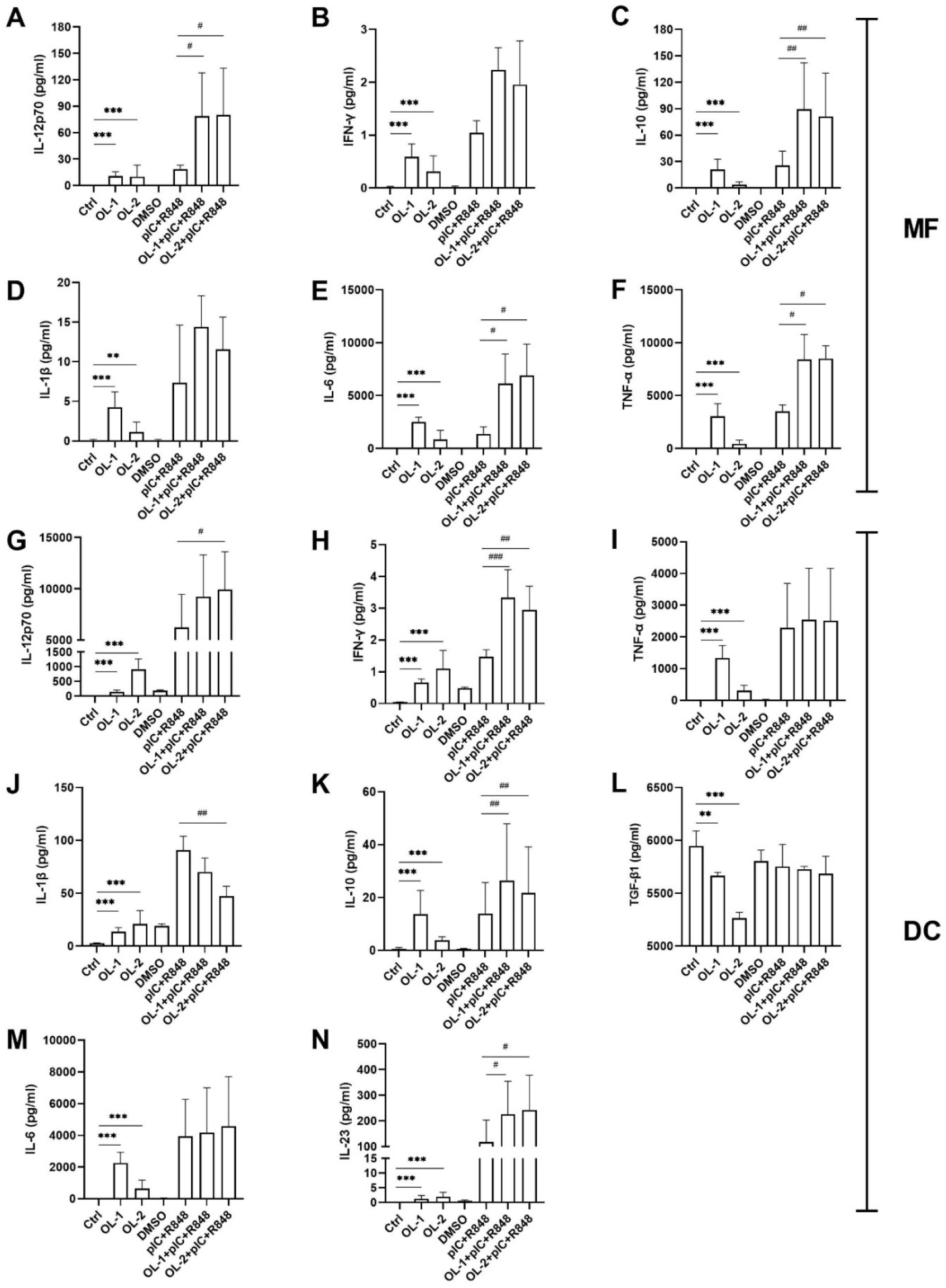
The effect of the consortia and pIC+R848 on human macrophage and dendritic cell cytokine response. Mfs and DCs were incubated with the treatments, for 24 and 48 hrs, respectively. Cytokine concentrations in the supernatants were measured with ELISA. DMSO dimethyl sulfoxide. The data was analyzed with linear model using change from a baseline as response (Ctrl vs OL-1 or OL-2; or pIC+R848 vs OL-1+pIC+R848 or OL-2+pIC+R848). N=4 * p=0.01 =< 0.05; ** p=0.001 =< 0.01; *** p<0.001 vs ctrl and # p=0.01 =< 0.05; ## p=0.001 =< 0.01; ### p<0.001 vs pIC+ R848.

Incubation of the OL-1 or OL-2 with Mfs for 24 hrs resulted in an increase in all the measured cytokines from supernatant by ELISA of which IL-6 and TNF-α had most pronounced increase, followed by IL-10, IL-12p70, IFN-γ, and IL-1β (**Figure 4A-4F**). Increase in cytokine production and type I immunity cytokines indicate Mf activation by both consortia. Modulation of pIC+R848 response by OL-1 and OL-2 resulted in an enhanced response to IL-6, IL-10, IL-12p70, and TNF-α (**Figure 4A-4F**), indicating additive stimulation of both pro-and anti-inflammatory responses during TLR3 and TLR7/8 stimulation.

After 48 hrs of incubation with DCs both consortia increased secretion of IL-12p70 compared to control (**Figure 4G**) and IFN-γ with or without pIC+R848 (**Figure 4H**), indicating Th1 polarization of the DCs by both consortia directly, but also augmentation of Th1 response upon TLR3 and TLR7/8 stimulation. TNF-α and IL-1β may support Th1 polarization of DCs, however, the effects of the consortia on the cytokine production were mixed as direct incubation increased TNF-α and IL-1β, but in the pIC+R848 response modulation there was either no effect or decrease of IL-1β in OL-2 samples (**Figure 4I and 4J**). Both consortia increased IL-10 production (**Figure 4K**) but suppressed TGF-β (**Figure 4L**), supporting polarization towards Th1, although some potentiation of the Th17 polarizing cytokines IL-23 and IL-6 secretion was observed (**Figure 4M and 4N**). In summary, the results suggest that consortia could drive polarization of the DCs towards Th1 cell induction, but with some induction of Treg and Th17 associated cytokines.

#### Probiotic consortia induce innate immune transcriptomes in macrophages and dendritic cells

To further test the effect of OL-1 and OL-2 on immune cell stimulation and to compare to pIC+R848 response, we conducted transcriptomics analyses. For sequencing, RNA was extracted from the OL-1, OL-2, and pIC+R848 -stimulated Mfs and DCs from the experiments presented above (**Figure 4**).

In differential gene expression (DEG) analysis, OL-1 and OL-2 were contrasted to control and pIC+R848 to DMSO. In Mfs OL-1, OL-2, and pIC+R848 treatments resulted in 158, 523, and 1264 DEGs, respectively, whereas in the DCs OL-1, OL-2, and pIC+R848 treatments resulted in 501, 1264, 670 DEGs, respectively (**Supplemental Table 1. Figure 5**). The number of shared DEGs between OL-1, OL-2 and pIC+R848 was 1350 for Mfs and 596 for DCs. OL-1 and OL-2 treatments shared 308 DEGs in DCs and 184 in Mfs, with pIC+R848 having 1904 unique DEGs in DCs and 184 in Mfs. Overall, the results support the observed increase in cytokine production and cell stimulation by OL-1 and OL-2, and similarity with pIC+R848 stimulation (**Figure 4**).

**Figure 5.**
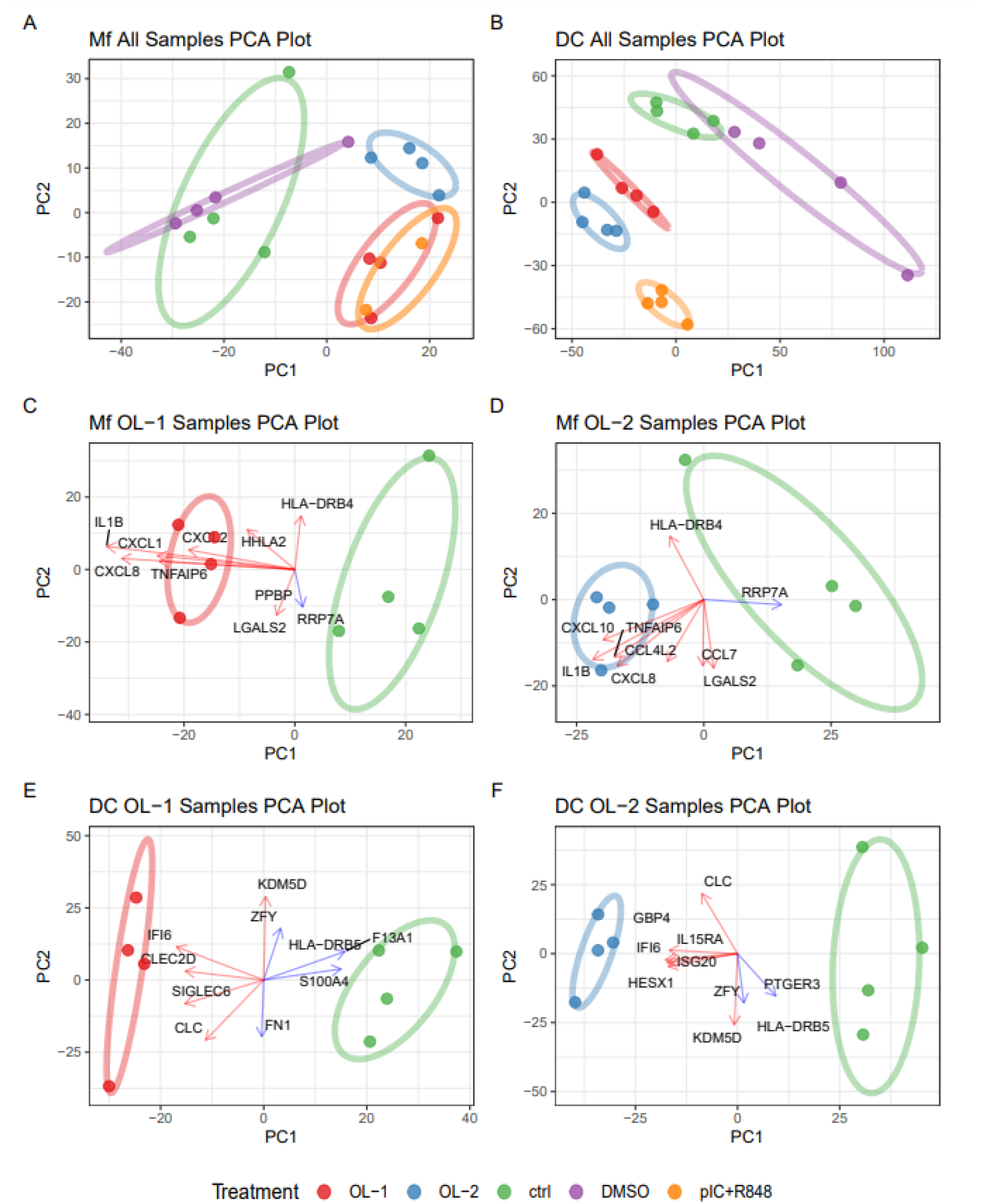
Global transcriptomics analyses of OL-1, OL-2, and pIC+R848 stimulated macrophages and dendritic cells. Principal component analysis (PCA) of transcriptomes of all samples tested for Mf (**A**) and DC (B). PCA plots including top gene loadings for Mf samples compared to the control for OL-1 (C) and OL-2 (E) and for Mf cells OL-1 (D) and DC (F). Color of the individual points and collective ellipses denotes treatment. Each point is an individual sample transcriptome containing all genes expressed. PC loadings are shown for OL-1 and OL-2 alone samples as arrows showing the effect of specific genes on the variation of the PCA plot. Arrows denote directionality of the variation and the value represents a graphical representation of the variation (in scale relation to each other, but not number on the scale) in either PC1 (horizontal axis) or PC2 (vertical). Color of the arrows represents the gene being upregulated (red) or down regulated (blue) in OL-1 or OL-2 vs ctrl comparison.

To have a broader view of the transcriptome similarities, we conducted a principal component analysis (PCA). Using both DC and Mf samples, PC1 captured 70% of the variance and PC2 14%, accounting for differences in the cell type and the treatment effect, respectively. The analysis showed that the blood cell donor added variation into the PCA, but was outweighed by the treatment effects. Due to differences in the transcriptomes of the cell types, DC and Mf datasets were analyzed separately. The PCA of the Mf data resulted in the separation of OL-1, OL-2, and pIC+R848 from the controls (DMSO and ctrl, PC1 49%), with OL-1 clustering closer to the pIC+R848 group than OL-2 (**Figure 5A**). The PCA of DC data showed a similar pattern (**Figure 5B**), however, in the DC data, PC1 accounted for a higher amount of variance than in Mf (68% vs 49%), suggesting a larger treatment effect on the transcriptome. In addition, pIC+R848 clustered separately from the control and the probiotic consortia. Overall, in Mf and DC datasets OL-1 and OL-2 clustered with pIC+R848 and separate from the controls, indicating potentially general similarity in immune gene activation. PC loadings for OL-1 Mf samples, showed major influence by *IL1B, CXCL8, CXCL1, CXCL2, PPBP* (CXCL7), and *TNFAIP6* on the PC1 (**Figure 5C**). Of these *IL1B, CXCL8*, and *TNFAIP6* were shared with OL-2, whereas *CCL4L2* and *CXCL10* were unique (**Figure 5D**). The results suggest that the key effects of the consortia versus control are on early innate and chemokine response. For DCs, OL-1 PCA showed separation of OL-1 from the control on PC1 with *IFI6, CLEC2D, SIGLEC6*, and *CLC* driving the effect (**Figure 5E**). Of these IFI6 and CLC were shared with OL-2, whereas *IL15RA, HESX1, ISG20*, and *GBP4* were unique (**Figure 5F**). *GBP4, IFI6, ISG20*, and *CLEC2D* genes are inducible by IFNs, and further CLC (Galectin-10) and CLEC2D (LLT1) expression by DCs are associated with modulating B and T cell responses.

We further investigated the effect of the OL-1 and OL-2 on the cytokine, chemokine, and co-stimulatory molecule gene expression. Overall, the profiles of OL-1 and OL-2 were similar, except for downregulation (*e*.*g. IL2, IL7, IL27, IL36G*) or upregulation (*e*.*g. IL6, IL32, TNFA*) of some transcripts in Mfs by OL-2 (**Figure 6**). Specifically for Mfs, OL-1 and OL-2 stimulation upregulated most prominently *EBI3* (IL27B), *IL1A, IL1B, IL12B*, and *IL23A*, necessary for type I immunity. Accordingly, chemokines profiles in Mfs showed broad activation of CCL and CXCL class chemokines, especially CCL1, CCL3, CCL4, CCL15, CXCL1, CXCL2, CXCL8, and CXCL10 that target neutrophils, Mfs, and NK cells. In addition, OL-1 and OL-2 induced HLA and co-stimulatory molecule expression, in line with innate response stimulation, but also upregulated inhibitory CD274 (Figure 6). In DCs, OL-1 and OL-2 induced upregulation of *IL2, IL15, IL18*, and *IL23* that are associated with T cell survival. Furthermore, treatments increased chemokine CX3CL1, CXCL9, CXCL10, CXCL11, CCL17, and CCL22 expression that are associated with attraction and activation of T cells. Probiotic consortia further inhibited CCL2, CCL3, and CCL4 expression that attract CCR2 and CCR5 positive cells such as innate NK cells and monocytes. Co-stimulatory gene expression results showed that DCs upregulated co-stimulatory genes CD80, CD83, CD86 after OL-1 or OL-2 treatment. Interestingly, inhibitory PD-ligands (CD274, PDCD1LG2) were upregulated but on the other hand tolerogenic response -associated CD31 (Clement et al., 2014) and CD36 (Lee et al., 2021) were downregulated. DCs showed upregulation of HLA class I and downregulation of HLA class II genes (**Figure 6**).

**Figure 6.**
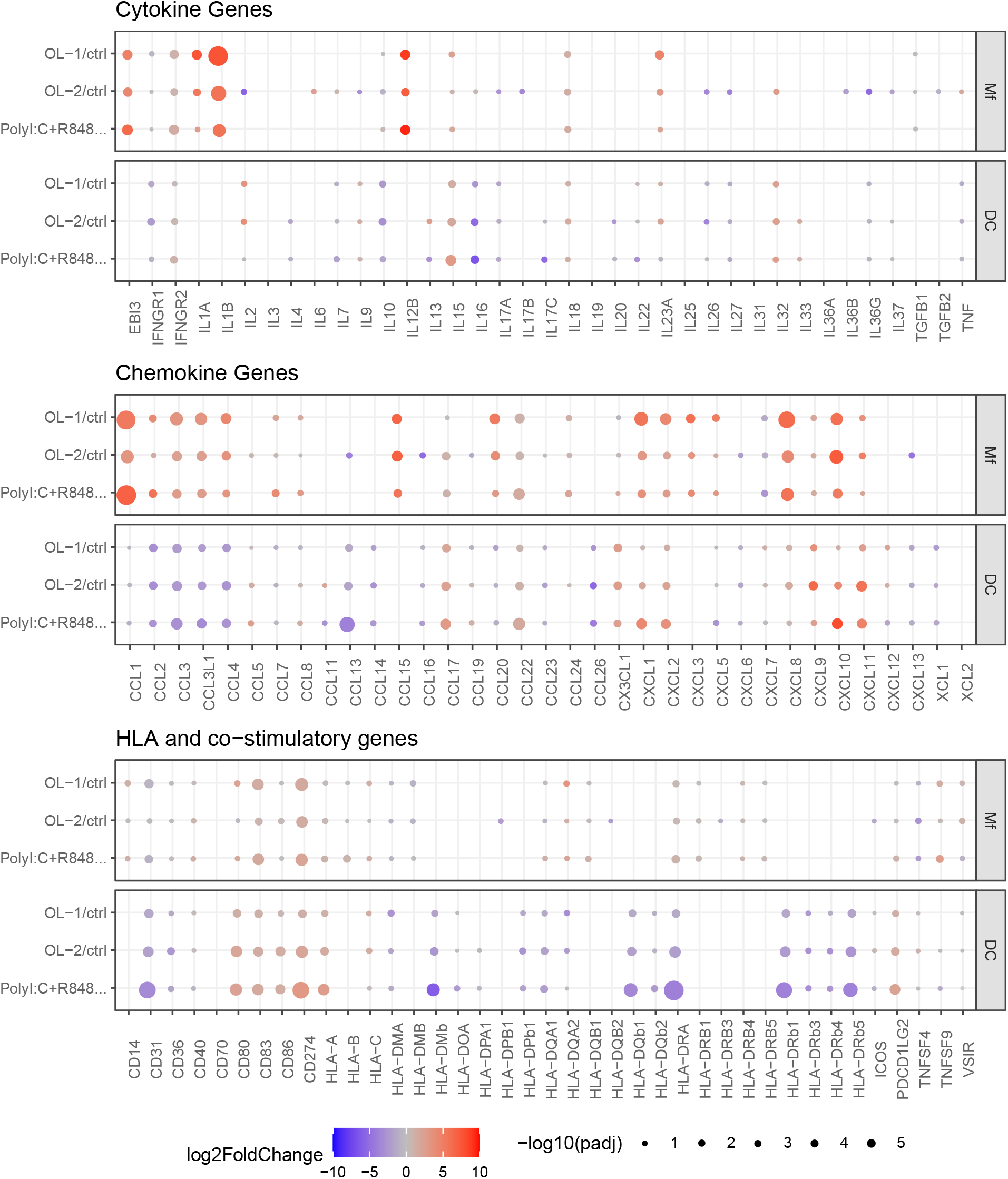
The effect of the OL-1, OL-2 and pIC+R848 on selected cytokines, chemokines and HLA/co-stimulatory gene response. Balloon plot of selected genes of importance from cytokines, chemokines and HLA/co-stimulatory gene groups. Only genes that show significant expression (padj <0.10) in either Mf or DC are shown. Size of the balloon denotes significance (-log10 adjusted p-value) with the larger size having more significance. Color denotes expression level (log2 fold change) with blue having reduced expression compared to the control and red having increased. Cell type is split in each panel denoted by Mf and DC.

#### OL-1 and OL-2 perturb COVID-19 KEGG pathway in human macrophages and dendritic cells

SARS-CoV-2 is known to suppress host cells anti-viral mechanisms to enable replication. As we had observed immune stimulation by OL-1 and OL-2, we wanted to further understand the effect of the consortia to anti-viral pathways critical for SARS-CoV-2 defense. We conducted pathway analysis using differential expression data which takes into account log2 fold changes and p-values for all genes in our dataset and calculates a probability of a KEGG pathway to be perturbed. Overall, OL-1, OL-2, and pIC+R848 affected 81, 36, and, 87 pathways in the Mfs and 71, 79, and 105 pathways in DCs, respectively (adjusted p-value < 0.1, FDR) (**Supplemental Table 2. Figure 7**). Pathways involving viral infections, microbial recognition, and innate and adaptive immune cell signaling pathway activation were broadly perturbed (**Supplemental Table 2. Figure 7**), indicating a general and broad impact by both OL-1 and OL-2 on key human immune defense pathways. In Mfs, OL-2 stimulated less pathways than OL-1, indicating perhaps milder stimulation that would be in line with lower cytokine concentrations in Mfs (**Figure 4**).

**Figure 7.**
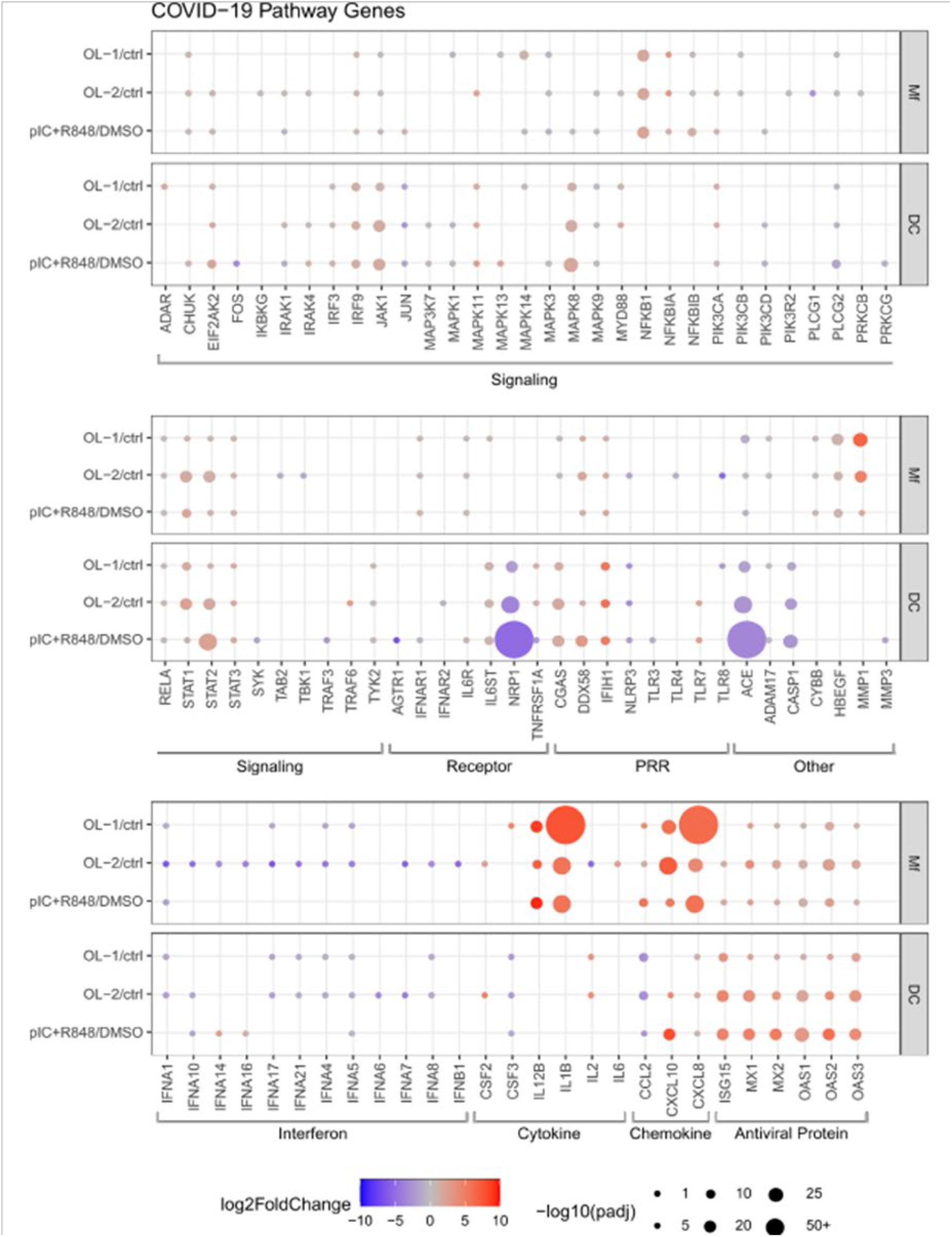
COVID-19 pathway gene expression for OL-1, OL-2 and plC+R848. Balloon plot of expression of genes taken from the Coronavirus disease – COVID-19 pathway in both Mf and DC cells. Only genes that have an adjusted p-value < 0.1 (-log10 >1) in either Mf or DC are shown. Size of the balloon denotes the log base 10 of the adjusted p-value, with smaller p-values having larger balloons. Color denotes expression level with blue having reduced expression compared to the control and red having increased. Black bars under the genes denote the function of the set of genes within the pathway. Cell type is split in each panel denoted by Mf and DC.

The results of the pathway analysis showed that OL-1 and pIC+R848 resulted in a significant perturbation of the COVID-19 KEGG pathway (KEGG: hsa05171), in Mfs (p = 0.03 (OL-1), p = 0.10 (OL-2), p = 0.35 (pIC+R848) and DCs, (p= 0.23 (OL-1), p = 0.15 (OL-2), p = 0.0005 (pIC+R848)), respectively (**Figure 7**). The “Coronavirus disease – COVID-19” KEGG pathway consists of 205 genes and includes all stages of viral infection, replication, release, and the SARS-CoV-2 induced innate immune and complement activation. In general, the gene expression pattern showed cell type specific gene expression regulation between Mf and DC after OL-1, OL-2, and plC+R848 stimulation (**Figure 7**).

In Mfs and in response to OL-1 and OL-2, interleukin IL12B and IL1B and chemokine CXCL10 and CXCL8 genes were highly expressed compared to low or no induction in DCs (**Figure 7**). However, the antiviral genes *MX1* and *MX2, OAS1, OAS2, OAS3* and *ISG15* were strongly induced in DCs and to a somewhat lower extent in Mfs. Interestingly, in DCs the neuropilin 1 receptor gene NRP1 involved in SARS-CoV-2 entry (Cantuti-Castelvetri et al., 2020) was reduced by all the treatments, as well as the gene coding ACE, whereas IFIH1 gene coding for MDA5 receptor was upregulated. Monocyte-derived innate immune cells seem to express the genes for SARS-CoV-2 receptor ACE2 and TMPRSS2 protease (Yang et al., 2020) that is needed for the activation of the viral spike protein and facilitation of the virus entry into the cell, but we did not observe these genes to be regulated by the treatments. Matrix metalloproteinase 1 gene (*MMP1*) expression was seen only in Mfs and the expression was higher in OL-1 stimulated cells compared to the others. Many of the JAK-STAT signaling pathway genes were upregulated (**Figure 7**), especially in DCs, indicating that the probiotic and plC+R848 activate IFN and ISG production in these cells.

## DISCUSSION

The potential of probiotics to support immune function against SARS-CoV-2 is yet to be established. In this study we have shown the effect of two probiotic consortia OL-1 and OL-2 on reducing nasal wash viral load in a ferret SARS-CoV-2 challenge model. The consortia modulated cytokine, chemokine, and IFN gene expression profiles in lungs and duodenum suggesting innate immune stimulation by OL-1 and OL-2. In human monocyte-derived Mf and DC model we further show that both consortia stimulate Th1 type cytokines and activate IFN response transcriptomic pathways, critical for innate defense against SARS-CoV-2. Further, the data suggests that the consortia modulates main receptor ACE2 in ferrets and may modulate co-receptors for viral entry in humans.

### Probiotics decrease SARS-CoV-2 nasal wash viral load

We evaluated the effect of OL-1 and OL-2 consortia in a pilot and main study on SARS-CoV-2 challenge. The results show a decrease in nasal wash viral load in the main study by both OL-1 and OL-2 probiotic consortia (Figure 1). In the main study, the time course and the magnitude of the SARS-CoV-2 viral titers in the placebo group was comparable to a recent SARS-CoV-2 dose response study in ferrets (Ryan et al., 2021), where the authors were using high (5 × 10^6^ plaque forming units (pfu)/ml), medium (5 × 10^4^ pfu/ml) and low challenge dose (5 × 10^2^ pfu/ml), of which the medium dose is comparable to 10^5^ TCID50 target dose in our study. Ryan et al showed around 10^7^ viral RNA copies/ml in the peak replication at 3 days with the high dose, whereas our lower viral dose resulted in around 10^7^ genomes/ml in the placebo group. They further showed that the initial viral inoculum plays a role in subsequent viral load. This indicates potential issues in the viral inoculum of our pilot study that was hampered by low infectivity of ferrets and low viral titers (**Figure S1**).

Although ferrets have been recognized as a good model for coronavirus infections, there are very few studies in ferrets where probiotics have been evaluated. Recently, a study assessed the effect of a probiotic consortia in ferrets and showed results on behavioral phenotypes after maternal pIC immune activation (Dugyala et al., 2020). However in mice models, probiotics administration has been shown to reduce viral load against influenza and some other respiratory pathogens (Lehtoranta et al., 2020) and in humans B. lactis Bl-04 (in consortia OL-1) supplementation was shown to decrease nasal wash viral titers in a rhinovirus challenge model (Turner et al., 2017) and risk of colds in healthy active adults (West et al., 2014). Our study results support the use of ferrets for evaluating the probiotic or microbial therapies efficacy against viral infections.

### Probiotic consortia modulate duodenal ACE2 and immune function

ACE2 is the main receptor for the SARS CoV-2 and it has been suggested that decreasing expression could limit viral entry into the cells, but studies on intestinal ACE2 are limited. It has been postulated that there is a potential intestinal infection and fecal–oral transmission of SARS-CoV-2 (Guo et al., 2021). In gut, ACE2 acts as a key regulator of dietary amino acid homeostasis, innate immunity, gut microbial ecology, and susceptibility to develop inflammation in a RAS independent manner (Hashimoto et al., 2012). In the current study on D5, during the acute infection, we observed by immunohistology a significantly higher expression of ACE2 in the duodenums of OL-1 and OL-2 group ferrets compared to placebo group animals (**Figure 3**), but by qPCR lower expression of ACE2 in all groups compared to D0 (**Figure 2**). As it has been shown that SARS-CoV-2 infection decreases ACE2 on the cell membrane (Verdecchia et al., 2020), the results suggest that OL-1 and OL-2 groups may have had lower duodenal viral infection and replication compared to placebo. This hypothesis is supported by increased *IFNA* expression by OL-1 and OL-2 compared to placebo at D5 (**Figure 2**) and suggests activation of the type I IFN pathway in the gut. On the other hand, ACE2 also acts as the SARS-CoV-2 receptor, and further, higher ACE2 expression has cytoprotective effects (Gheblawi et al., 2020), and thus it is not clear what is an optimal level of ACE2 expression (Chaudhry et al., 2020). Despite increase in ACE2, we did not observe differences in the duodenal tissue inflammation (**Figure 2**) or in inflammatory gene expression response between the treatments at D5 (**Figure 3**). Interestingly, the Mf and DC treatment with OL-1 and OL-2 decreased expression of ACE (**Figure 7**) that has opposite function to *ACE2*. The results indicate that the effect of the probiotic consortia on the gut ACE-ACE2 balance and RAS system could be cytoprotective. The transcriptomics data also showed a decreased expression of *NRP1* that acts as a cofactor for ACE2-mediated viral cell entry (**Figure 7**) (Cantuti-Castelvetri et al., 2020), which adds to the evidence on the influence of the OL-1 and OL-2 on SARS-CoV-2 receptors *in vivo* and *in vitro*.

At D10 in the main study, duodenal ACE2 expression decreased in OL-1 and OL-2 groups compared to placebo, and also compared to D5 OL-1 and OL-2 levels (**Figure 3**), perhaps suggesting faster resolution to baseline ACE2 expression in probiotic groups, however, with lack of non-infectious samples for baseline this hypothesis remains to be tested.

The inflammatory gene expression at D10 was comparable to D5 in all groups, however, the histological inflammatory scoring showed a decrease from D5 to D10 (**Figure 3I and 3J**), in line with resolution of intestinal infection. The effects of OL-1 and OL-2 compared to placebo were relatively pronounced and consistent with duodenal gene expression. Both consortia increased the expression of type III IFN *IFNL1*, monocyte (CCL2) and neutrophil (IL8) targeting chemokines, and *TLR8* that recognizes SARS-CoV-2, suggesting stimulation of anti-SARS-CoV-2 immunity. OL-2 also induced cytosolic SARS-CoV-2 pattern recognition receptor *IFIH1* (MDA5), *IL6*, and *TNFA*, and OL-1 decreased *IFNG*, that points to consortia specific differences on immune stimulation. The differences between OL-1 and OL-2 were also observed in duodenal gene expression in the pilot study, where OL-2 decreased IFN responses at D5, and OL-1 decreased pro-inflammatory and IFN gene expression at D21 compared to placebo. In addition, OL-1 and OL-2 showed significant reduction in ACE2 transcripts when compared to placebo at D21.

So far little has been done to evaluate therapies or intervention to target the gastrointestinal tract pool of SARS-CoV-2. Based on the differential expression of ACE2 in the duodenum we propose that OL-1 and OL-2 intervention could potentially be effective in managing the SARS-CoV-2 load in the duodenum, potentially also in humans as supported by decreased expression of NRP1 and ACE in human DC model. This is a first report to show probiotic-mediated changes in ACE2 expression in a SARS-CoV-2 infection model.

### Probiotic delivery by gavage may stimulate gut-lung immune axis

In COVID-19, lungs are the key site for pathology and viral replication. The gene expression results at D5 and D10 of the main study compared to non-infected ferrets from the pilot study, show that SARS-CoV-2 induces type I (*IFNA*) and III (*IFNL1*) IFN responses, but suppresses type II response (*IFNG*), and stimulates pro-inflammatory (*IL6 and IL8*) and anti-inflammatory (*IL10*) gene expression (**Figure 2**). Overall, the cytokine profiles indicate balanced response of the ferrets to the virus where the viral replication is controlled without cytokine storm. We also observed at D5 and D10 in all groups increased expression of *ACE2* that has anti-inflammatory and cytoprotective properties in addition to its role as a receptor for SARS-CoV-2 (Gheblawi et al., 2020). The results above are in line with the lack of symptoms in the ferrets and low viral load by D10 (**Figure 1**). In the pilot study minimal changes in gene expression was observed that likely reflects the low infectivity (**Figure S1**). Similar immune responses in ferrets to SARS-CoV-2 have been observed previously (Blanco-Melo et al., 2020) that supports data from our unconventional use of non-infected animals from the pilot study to normalize the response of the infection in the main study, however, the results need to be interpreted cautiously.

Probiotic effect compared to placebo on the lung gene expression was low. In the main study at D5, OL-2 increased expression of chemokines (*CXCL10 and IL8*) targeting NK cells and neutrophils, and *TLR8* that recognizes viral RNA (**Figure 2**). This effect was not observed at D10 and could be due to resolution of the infection. In the pilot study OL-1 at D0 decreased *CXCL10* and *TLR8*, suggesting effects on the lung immune function pre-infection, however, no changes were observed in duodenum. At D5 OL-1 increased *IFNG* and at D21 post infection the differences were most pronounced for OL-1 that increased *IFNA, IFNL1, IL10, IL6*, and *TNFA*, and decreased *CCL2*, also suggesting effects post-infection. In summary, probiotic consortia seem to induce lung immune system, but the effects seem mild and somewhat inconsistent in this data set. On the other hand, OL-1 and OL-2 reduced the viral load that is suggestive of influence on nasal immune system as well. The effect of the probiotic consortia on gut-lung or gut-respiratory axis warrants further investigation.

### OL-1 and OL-2 may prime innate immune responses against viral infections

Innate immune systems between ferrets and humans, and mammals in general are homologous, however, to gain further understanding on potential effect of the consortia in humans we exposed human blood-derived Mfs and DCs to OL-1 and OL-2 and viral RNA analog cocktail pIC+R848 and analyzed their cytokine and transcriptome responses.

The studies on SARS-CoV-2 infection and pathogenesis indicate that inadequate innate immune and IFN response increase susceptibility to more severe COVID-19 (Schultze and Aschenbrenner, 2021) that is supported by increased risk of more severe disease by aging. Our results show that exposure of Mfs to OL-1 and OL-2 resulted in the secretion of cytokines important for Mf activation and induction of type I immunity (IL-12 and IFN-γ), but also drove pro-and anti-inflammatory cytokine production (**Figure 4**). These results were supported by transcriptomics analysis showing activation of IL-1 and IL-12 cytokine family genes, upregulation of co-stimulatory molecules, MHC-I/II and chemokines targeting monocytes, NK cells, and neutrophils critical for early defense in viral infections (**Figure 6**). DC response to OL-1 and OL-2 further show Th1 polarization as determined by increased IL-12p70 production (**Figure 4**). The transcriptomics analysis further showed an increase in STAT1 (**Figure 7**) that drives in DCs Th1 polarization (Johnson and Scott, 2007), induction of co-stimulatory and MHC-I molecules, and upregulation of genes related to T cell growth and attraction. Lu et al. 2021 showed that SARS CoV-2 (24h infection with clinical isolates) induces *IL1A, IL1B, IL8, CXCL10, CCL2*, and *CCL3* mRNA in human PBMC-derived myeloid cells (Lu et al., 2021). We see quite similar gene induction pattern in our monocyte-derived Mfs (**Figure 7**), where both consortia and pIC+R848 induce some of the same genes as SARS-CoV-2, suggesting induction of anti-viral pathways. In summary, the results show that probiotic consortia could prime or train the innate immune system also in humans prior to SARS-CoV-2 infection to enhance resilience of the host.

Priming of the innate system by OL-1 or OL-2 is a potential mechanism to explain the reduced viral load that we observed in the ferrets after 7 days of supplementation (**Figure 1**). Although we have limited data pre-infection, the post infection gene expression results suggest enhanced response in duodenum, but also potentially in lungs (**Figure 2**). Netea et al. have suggested that the training of the human innate immune system, for example by vaccines, prior to SARS-CoV-2 infection could reduce the susceptibility to and the risk of severe COVID-19 (Netea et al., 2020). Similar evidence on priming of the immune system against viral infections by probiotics have been noted more broadly in vitro and in human clinicals (Lehtoranta et al., 2020; Turner et al., 2017).

### Probiotic consortia may counteract SARS-CoV-2 immune-evasion

Studies on the early pathogenesis of SARS-CoV-2 show that it infects the cells via ACE2 that is internalized. The host cells recognize the virus by endosomal (TLR3, TLR7, and TLR8) and cytosolic (RIG-I and MDA5) receptors that recognize ssRNA and dsRNA. This leads to an MyD88 and NF-κB -dependent activation of pro-inflammatory response and IRF3/7 -dependent type I IFN response. IFNs drive STAT1/2 and IRF9 mediated ISG response. SARS-CoV-2 produces proteins that suppress the activation of this cascade (Schultze and Aschenbrenner, 2021). The results of our transcriptomics study show that OL-1 can significantly modulate the KEGG COVID-19 pathway in Mfs, and that OL-2 shows similar gene expression pattern in Mfs but does not result in a significant change on a pathway level (**Figure 7**). In DCs, the result on a pathway level are significant only for pIC+R848, however, OL-1 and OL-2 show similar pattern of significant gene expression (**Figure 7**). Several signaling molecules on the pathways in Mfs and DCs show upregulation by the consortia including *IRF3, IRF9, NFKB1, STAT1, STAT2*, and *ISGs ISG15, MX1, MX2, OAS1, OAS2, OAS3*, and *CGAS*. In addition, viral RNA sensors *IFIH1* (MDA5) and DDX58 (RIG-I) show an increased expression, whereas *TLR7* and *TLR8* expression have mixed results. Interestingly type I IFN family gene expression seems to be suppressed or unimpacted by OL-1 and OL-2, but mainly also by pIC+R848 (**Figure 7**). Overall, the results suggest that OL-1 especially, but also OL-2, could potentially counteract suppression of the innate immune system activation by SARS-CoV-2. Delayed or poor IFN response has been associated with more severe COVID-19 disease in human clinical samples (Hadjadj et al., 2020) and lower levels of pro-inflammatory cytokines and chemokines noted with SARS-CoV-2 infection (Blanco-Melo et al., 2020; Chu et al., 2020), suggesting that targeting innate immunity with probiotic consortia could provide benefits in managing SARS-CoV-2 infection in humans.

## Conclusions

In this study we have shown that probiotic consortia OL-1 and OL-2 stimulate ferret immune function and reduce viral load during SARS-CoV-2 infection (**Figure 8**). Results of the human immune cell stimulation indicate that pathways and cytokine secretion associated with SARS-CoV-2 immune defense are activated and thus these probiotic consortia could potentially provide benefits to support immune function against SARS-CoV-2, perhaps by priming or training the innate immune system (Lehtoranta et al., 2020; Netea et al., 2020) (**Figure 8**). We have also shown influence of the probiotic consortia on duodenal immune function and ACE2 expression. As many COVID-19 patients suffer from gastrointestinal symptoms, these probiotic consortia could potentially support intestinal health and immune function. Probiotics in general have shown efficacy in meta-analyses against respiratory tract infections (Hao et al., 2015; King et al., 2014), however, the results between the strains or their combinations vary. Thus, it is warranted to use specific strains or consortia of probiotics for immune stimulation (Hill et al., 2014). Even small differences in genome or within strains may result in vast differences in phenotypes and metabolism (Morovic et al., 2018; Zabel et al., 2020). Human clinical studies should be urgently initiated to better understand the effect of OL-1 and OL-2, and other microbial therapies in managing SARS-CoV-2 infections.

**Figure 8.**
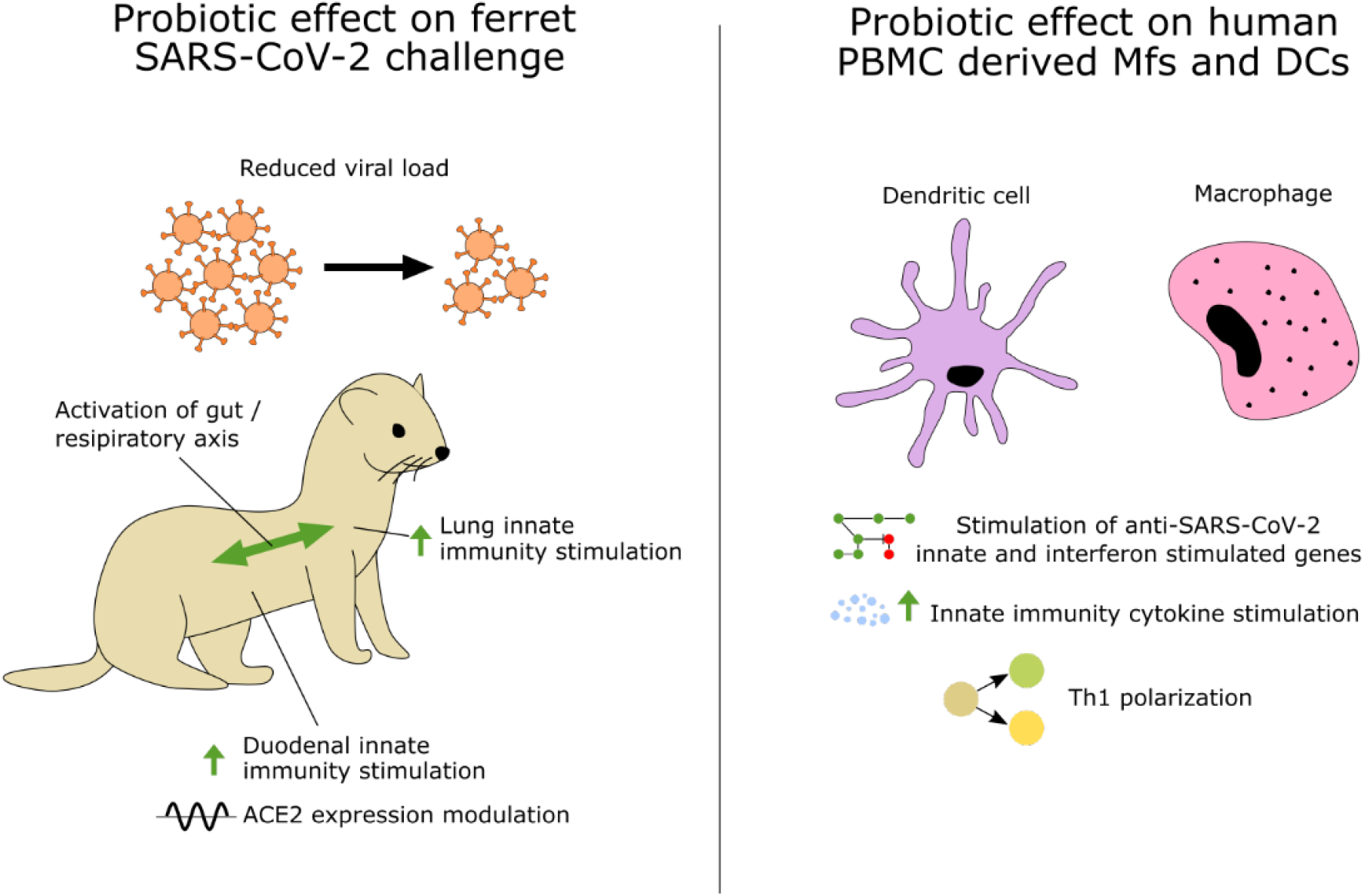
Summary of the effect of the probiotic consortia in ferrets on SARS-CoV-2 challenge and human Mfs and DCs. Probiotic gavage of OL-1 and OL-2 stimulate ferret duodenal and respiratory tract innate immunity and reduce SARS-CoV-2 viral load in nasal washes, thus suggesting activation of gut-respiratory axis immunity. Further, Mf and DC stimulation by OL-1 and OL-2 show activation of critical genes and secretion of cytokines important for anti-SARS-CoV-2 innate immunity, suggesting that these probiotic consortia could also be beneficial in humans.

## ACKNOWLEDGMENTS

Jaana Larsson-Leskelä, Katri Holappa, Henri Ahokoski, Anna Lappalainen for excellent technical work, and Ilmari Ahonen for statistical analysis of the ELISA data. Xinjian Peng, PhD; Senior Biologist, IIT Research Institute (IITRI)

## COMPETING INTERESTS

All authors are employees of IFF that manufactures and sells probiotics used in the study.

## Materials and Methods

### Probiotic strains and placebo

The probiotic strains used in the experiments were for OL-1: *Bifidobacterium longum* subsp. *infantis* Bi-26, *Danisco* Global Culture Collection (DGCC)11473, *Bifidobacterium animalis* subsp. *lactis* Bl-04, American Type Culture Collection (ATCC) Safe Deposit (SD)5219, DGCC2908, *Lacticaseibacillus paracasei* subsp. *paracasei* Lpc-37, SD5275, ATCC PTA-4798, *Lacticaseibacillus rhamnosus* Lr-32, SD5217, and *Ligilactobacillus salivarius* Ls-33, SD5208, and for OL-2: *Bifidobacterium animalis* subsp. *lactis* Bi-07, SD5220, DGCC 2907, Lactobacillus acidophilus NCFM, SD5221, ATCC 700396, *Limosilactobacillu*s *fermentum* SBS-1, DGCC1925, *Lactococcus lactis* subsp. *lactis* Ll-23, DGCC8656, Streptococcus *thermophilus* subsp. *thermophilus* St-21, SD5207. Potato *maltodextrin* was used as a placebo. All the strains are owned by and were manufactured along with placebo by IFF (Madison, Wisconsin, USA)

### Ferret studies

Outbred male ferrets (Triple F Farms, Sayre, PA) were quarantined at IIT Research Institute (IITRI), Chicago, USA for approximately 6-7 days before initiation of the dosing. Animals were approximately 8-11 months of age and were approximately 1.0 to 2.0 kg at the initiation of dosing. Animals were maintained in BSL-2 facility and were transferred to BSL-3 before infection. Ferrets were randomly assigned to the different experimental groups.

Pilot Study: From D-21 through study D21, ferrets in group 1 received a daily oral gavage with the OL-1 consortia (20B CFU/ strain), and ferrets in group 3 received a daily oral gavage with the Placebo. Ferrets in group 2 received a daily oral gavage with the OL-2 consortia (20B CFU/ strain) from study D1 through study D21. The virus was administered intranasally (i.n.) as droplets on study D0. On study D1, D3, D5, D7, and D9 nasal washes were collected. Five ferrets from each OL-1 and placebo groups were necropsied on D0 and five ferrets from each group were necropsied on D5 and D21 in the pilot study.

Main Study: From study D-7 through study D10, ferrets in group 1 received a daily oral gavage with the OL-1 consortia (20B CFU/ strain), ferrets in group 2 received a daily oral gavage with the OL-2 consortia (20B CFU/ strain), and ferrets in group 3 received a daily oral gavage with the Placebo. Each probiotic capsule was opened and content diluted in 3 mL sterile PBS for each daily oral gavage. 2019 Novel Coronavirus, isolate USA-WA1/2020 (SARS-CoV-2) was used to challenge the ferrets. The viruses were produced by infecting VE6 cells at IITRI with the SARS-CoV-2 and tissue culture infection dose (TCID50) was calculated. The virus in DMEM was administered i.n. as droplets on study day 8. Ferrets were anesthetized with Ketamine/Xylazine for delivery of the virus. A total of 0.5 mL of virus (target of 10^5^ TCID50) was delivered to each study ferret. The 0.5 mL dose was split between each nostril (0.25 mL/nare). To confirm the inoculation titer of the virus, aliquots of the prepared virus solution were collected, the aliquots were stored at ≤ -65 °C, and a viral titer assay (TCID50) was performed. On study days D1, D3, D5, D7, D9, and D10 nasal washes were collected. Ferrets were anesthetized with Ketamine/Xylazine, and 0.5mL of sterile PBS containing penicillin (100 U/mL), streptomycin (100 µg/mL) and gentamicin (50 µg/mL) were injected into each nostril and collected in a specimen cup when expelled by the ferret. The recovered nasal wash was collected, volume recorded, and aliquoted. One aliquot (∼100 µL) was treated with DNA/RNA Shield (Zymo Research, Irvine, CA) and then stored at room temperature for determination of viral load by RT-qPCR. Five ferrets from each group were necropsied on D5, and remaining ferrets from each group were necropsied at D10 in the main study. Lungs were collected and 0.5-1.0 cm^3^ piece of lung was flash frozen, and then stored at ≤ -65 °C. The duodenum was harvested, cut longitudinally to expose the inner mucosal layer, rinsed with sterile PBS, and divided into five roughly equal portions. Samples were flash frozen and stored at ≤ -65 °C for RNA isolation and samples were fixed in 10% formaline for IHC.

The study was performed under American Association for Laboratory Animal Science (IACUC) protocol #20-031

### RT-qPCR SARS-CoV-2 Titers

The concentration of virus in nasal washes from study days D1, D3, D5, D7, D9, and D10 was determined by RT-qPCR assay. Briefly, RNA was extracted from samples stored in RNA/DNA Shield using the Quick-RNA Viral Kit (Zymo Research) according to manufacturer’s protocol. RNA was eluted with 100 µL nuclease-free water. A standard curve was prepared by using blank ferret nasal wash collected from ten naïve ferrets and spiked with known concentrations of viral RNA. Each RT-qPCR plate included 9 RNA standards (5 × 10^7^, 5 × 10^6^, 5 × 10^5^, 5 × 10^4^, 5 × 10^3^, 5 × 10^2^, 50, 20, 5 copies per RT-qPCR well) in duplicate, a NTC (no template control) and a positive control in triplicate well. Each test sample was analyzed in duplicate wells.

RT-qPCR was performed using the iTaq universal probes onestep kit (Bio-Rad). 5 µl viral RNA was used for RT-qPCR. Total reaction volume was 15 µL (5 µl RNA + 10 µL master mix). The following RT-PCR cycling conditions will be used: 50 °C for 15 min (RT), then 95 °C for 2 min (denature), then 40 cycles of 10s at 95 °C, 45s at 62 °C.

Primers used for SARS-CoV-2 detection:

2019-nCoV_N1-F 5’-GACCCCAAAATCAGCGAAAT-3’

2019-nCoV_N1-R 5’-TCTGGTTACTGCCAGTTGAATCTG -3’

Probe: 2019-nCoV_N1-P: 5’-FAM-ACCCCGCATTACGTTTGGTGGACC-BHQ1-3’

The area under the curve (AUC) over the 10 days post infection was calculated from the viral titer trend lines for each ferret. Kruskal-Wallis nonparametric test was performed to evaluate the differences in the AUC among the three treatment groups (sig. level = 0.05), and post-hoc pairwise Wilcoxon tests were deployed for multiple comparisons using Benjamini-Hochberg method for p-value adjustment.

### Gene expression analysis by qPCR

Lung and duodenum were collected and stored in trizol. Tissue RNA was isolated using Direct-Zol RNA Miniprep Plus kit. Two step RT qPCR were used to analyze the gene expression in ferret lung and duodenum total RNA samples. Reverse transcription was performed with iScript Reverse Transcription Supermix (Bio-Rad). For lungs, 1250 ng RNA per 50 µl RT reaction and for duodenum, 625 ng RNA per 50 µl RT reaction was used. The total RT reaction volume was 50 µL and RT was performed in a 96 well plate on a Bio-Rad CFX96 Touch Real Time PCR Detection System. After RT, the cDNA was diluted with nuclease free water to bring the final total volume up to 200 µL (4x dilution). 5 µL of diluted cDNA was used for qPCR analysis of the expression of gene of interest. The total PCR reaction volume was 15 µL per well (10 µL master mix + 5 µL cDNA sample) and the qPCR was performed on the CFX384 Touch Real-Time PCR Detection System (Bio-Rad) using the cycling conditions listed below:

| Temperature | Time | Cycle(s) |
| --- | --- | --- |
| 95 °C | 2 minutes | 1 |
| 95 °C | 10 seconds | 40 |
| *60 °C or 62 °C | 45 seconds |  |

\*60 °C for CCL2, IFN $\alpha$ , IFNL1, and IL8

62 °C for all other genes (CCL5, CXCL10, IFNγ, IL10, IL1β, IL6, TLR8, TNFα, ACE2, and RPS18)

Each plate contains a positive control (prepared from stimulated ferret PBMC cells) in duplicate wells, an NTC in duplicate wells and all test samples in duplicate wells. Primer and probe sequences can be found in Supplemental Table 3.

The threshold baseline of the qPCR assay was manually set at 200 for all plates if the automatically calculated (by the CFX Maestro 1.1 software) threshold baseline was >200. The 200-threshold baseline value was determined at the development/qualification stage of RT-qPCR assays of genes of interest. However, if the automatically calculated threshold baseline was < 200, then the threshold baseline was directly used. Cq value in each well was also automatically calculated by the Maestro 1.1 software based on the threshold baseline. Data generated by the software were exported to a Microsoft Excel (Microsoft Corporation; Redmond, WA) spreadsheet for data processing.

dCq was calculated by subtracting cytokine mRNA gene level from that of GAPDH. Then ddCq was obtained by subtracting dCq from placebo group results. The distribution of dCq (grouped by gene, tissue, and day) was then examined for normality (Shapiro test and qq plot) and extremes (< 1Q-3×IQR or > 3Q+3×IQR). After removal of extremes, mixed effects models were performed for each gene:

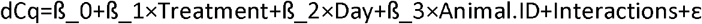

If the interaction effect Treatment×Day was significant, ANOVA was carried out to compare dCq across three treatment groups within the same tissue and day, followed by post-hoc analysis through obtaining estimated marginal means (EMMs) (Lenth et al., 2021) and pairwise comparison thereof. The multiple comparison p-values were adjusted by Benjamini-Hochberg (BH) method. Log2 fold change was calculated using the log2 of the ratio of EMM dCq test/EMM dCq control. For EMM dCq control we used the placebo for each gene at the same time point or the pre-infection D0 placebo time point for each gene tested.

### Immunohistochemical analysis of duodenum for ACE2 expression

Tissue samples were collected from PBS control and SARS CoV-2 -infected ferrets and incubated in 10% neutral-buffered formalin for fixation before they were embedded in paraffin based to standard procedures. The embedded tissues were sectioned and dried for 3 days at room temperature. To detect the ACE2 by immunohistochemistry, Goat Anti-ACE2 antibody (R&D, Cat#: AF933, Lot#: HOK0620051) was used as the primary antibody. Antigen was visualized using citrate buffer (pH6.0), microwaved for 3 min and steam for 15 min.

Slides were viewed and digitized pictures were taken by Olympus VS120 scanner and analyzed by a senior Pathologist using Olyvia 3.2 software (Olympus Corporation, Tokyo, Japan).

### Histopathological examination of duodenum

ACE2 expression was scored by using H-score. The H-score is based on a predominant staining intensity. The Intensity scores are classified as negative (0), low intensity (2), medium intensity (2) and high intensity (3). The percentage of cells at each staining intensity level is counted (estimated), and finally, an H-score is assigned using the following formula:

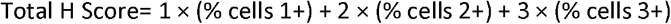

### PBMC stimulation assay

Probiotic strains were grown to logarithmic growth phase, collected by centrifugation, washed once with PBS and suspended to cell culture medium. The optical density (OD)600 was adjusted to correspond to bacteria:host cell ratio of 10:1. The bacteria in the consortia were applied on cells in equal proportions 2+2+2+2+2.

Monocytes were purified from freshly collected leukocyte-rich buffy coats obtained from healthy blood donors through the Finnish Red Cross Blood Service (permission no 46/2016, renewed annually). The use of human blood was approved by the Ethics Committee of the Hospital District of Helsinki and Uusimaa, Finland (216/13/03/00/2016). Human peripheral blood mononuclear cells were isolated by density gradient centrifugation followed by purification of monocytes with CD14+ magnetic beads. To obtain Mfs purified monocytes were plated on 24 well plates 3 × 10^5^ cells/well and cultured for 7 days in Macrophage-SFM (Gibco, Life Technologies, Grand Island, NY, USA) with recombinant human GM-CSF (Miltenyi Biotech) 1000 IU/ml and 1 % Antibiotic-Antimycotic. To differentiate monocytes into immature dendritic cells, monocytes were plated on 12 well plates 5 × 10^5^ cells/well (Falcon, Corning, NY, USA) and cultured for 7 d in RPMI-1640 (Sigma) supplemented with 1% Antibiotic-Antimycotic, 10% fetal bovine serum (FBS), IL-4 (400 IU/ml) and GM-CSF (1000 IU/ml).

Cells from four blood donors were stimulated with consortia alone (bacteria:cell ratio of 10:1) or in combination with TLR ligand blend, pIC, 30μg/ml + R848, 10μM (both from Sigma-Aldrich, St. Louis, MO, USA) for 24 h (Mf) or 48 h (DC). PBMCs without stimulation were used as a control, dimethyl sulfoxide (DMSO), a carrier for the TLR-ligands, were used as a control for pIC+R848 containing treatments.

### Enzyme linked immunosorbent assay

Cell culture supernatants from Mf and DC cultures are analyzed for IFN-γ, IL-1β, IL-6, IL-10, IL-12p70, and TNF-α by Quanterix Human 6-Plex Ciraplex Array (Quanterix, Inc., Billerica, MA, USA). In addition, DC supernatants were analyzed for IL-23 and TGF-β by Quanterix Human 1-plex Assays according to manufacturer’s instructions. Results were analyzed with CiraSoft software (Quanterix).

The data were log2-transformed prior to statistical analysis. The data were analyzed using a linear model that included the main effects of treatment and donor (2-way ANOVA). Pairwise-comparisons between the treatments were performed using contrasts of estimated marginal means and corrected for false positive rate using Sidak’s method. The fit of the models was checked by inspecting the normality of the model residuals. One extreme low value was removed from the analysis to meet model assumptions (DC: DMSO, IL-23).

### PBMC RNA extraction

Monocyte-derived macrophages and DCs were lysed upon collection with MagMAX(tm) Lysis/Binding Solution (Invitrogen, Thermo Fisher Scientific) and frozen in -80 °C. RNA was extracted with KingFisher Flex automated extraction instrument (Thermo Scientific, Thermo Fisher Scientific Inc. Waltham, MA, USA) using MagMAX(tm)-96 Total RNA Isolation Kit (Invitrogen, Thermo Fisher Scientific) following manufacturer’s instructions. The RNA quality and quantity were analyzed with Agilent 4200 TapeStation System (Agilent, Santa Clara, CA, USA).

### Transcriptomics

Targeted RNA-seq library preparation was conducted with Tempo-Seq (BioSpyder) detector oligonucleotides designed to target all human genes (∼23,000). Library preparation was conducted according to BioSpyder kit user manual. 40 Samples were pooled, purified, quantitated and stored frozen prior to sequencing. The pooled libraries were sent to Roy J. Carver Biotechnology Center at the University of Illinois at Urbana-Champaign for TempO-Seq sequencing using the Illumina NovaSeq platform (1×50nt). The target read-depth per sample was 5 million reads with the minimum reads per sample at 1 million. The reads were quality filtered, trimmed and demultiplexed and aligned to the BioSpyder Human Whole Transcriptome v2.0 annotated reference using Salmon (v1.1.0)(Patro et al., 2017). Two samples did not pass the 5 million read minimum, which were both from pIC+R848 treated DC and were not included in the analysis. 38 total samples were processed, sample quality metrics for RNA quality and alignment to the reference can be found in Supplemental Table 4.

Differential expression analysis was conducted using DESeq2 (v1.26.1)(Love et al., 2014) in R (v 3.6.2). Mf and DC were analyzed separately as initial analysis determined the cell types to have distinct expression profiles. Genes were considered differentially expressed if the adjusted p-value < 0.1 and the absolute value of the log2 fold change > 1. Principal component analysis (PCA) was performed to determine overall relatedness of the samples under the various experimental conditions using ggplot2 (v 3.3.3) (Wickham, 2016). Pathway analysis was performed using ROntoTools (v 2.14) (Voichita et al., 2019) with a KEGG pathway (Kanehisa, 2019; Kanehisa et al., 2021; Kanehisa and Goto, 2000) considered significantly changed with a false discovery rate perturbation value of > 0.1.

## REFERENCES

Alfi, O., Yakirevitch, A., Wald, O., Wandel, O., Izhar, U., Oiknine-Djian, E., Nevo, Y., Elgavish, S., Dagan, E., Madgar, O., et al. (2021). Human Nasal and Lung Tissues Infected Ex Vivo with SARS-CoV-2 Provide Insights into Differential Tissue-Specific and Virus-Specific Innate Immune Responses in the Upper and Lower Respiratory Tract. Journal of Virology 95, e00130–00121.

Blanco-Melo, D., Nilsson-Payant, B.E., Liu, W.C., Uhl, S., Hoagland, D., Moller, R., Jordan, T.X., Oishi, K., Panis, M., Sachs, D., et al. (2020). Imbalanced Host Response to SARS-CoV-2 Drives Development of COVID-19. Cell 181, 1036–1045 e1039.

Cantuti-Castelvetri, L., Ojha, R., Pedro, L.D., Djannatian, M., Franz, J., Kuivanen, S., van der Meer, F., Kallio, K., Kaya, T., Anastasina, M., et al. (2020). Neuropilin-1 facilitates SARS-CoV-2 cell entry and infectivity. Science 370, 856–860.

Chaudhry, F., Lavandero, S., Xie, X., Sabharwal, B., Zheng, Y.Y., Correa, A., Narula, J., and Levy, P. (2020). Manipulation of ACE2 expression in COVID-19. Open Heart 7.

Chu, H., Chan, J.F., Wang, Y., Yuen, T.T., Chai, Y., Hou, Y., Shuai, H., Yang, D., Hu, B., Huang, X., et al. (2020). Comparative Replication and Immune Activation Profiles of SARS-CoV-2 and SARS-CoV in Human Lungs: An Ex Vivo Study With Implications for the Pathogenesis of COVID-19. Clin Infect Dis 71, 1400–1409.

Clement, M., Fornasa, G., Guedj, K., Ben Mkaddem, S., Gaston, A.T., Khallou-Laschet, J., Morvan, M., Nicoletti, A., and Caligiuri, G. (2014). CD31 is a key coinhibitory receptor in the development of immunogenic dendritic cells. Proc Natl Acad Sci U S A 111, E1101–1110.

Dugyala, S., Ptacek, T.S., Simon, J.M., Li, Y., and Frohlich, F. (2020). Putative modulation of the gut microbiome by probiotics enhances preference for novelty in a preliminary double-blind placebo-controlled study in ferrets. Anim Microbiome 2.

Gheblawi, M., Wang, K., Viveiros, A., Nguyen, Q., Zhong, J.C., Turner, A.J., Raizada, M.K., Grant, M.B., and Oudit, G.Y. (2020). Angiotensin-Converting Enzyme 2: SARS-CoV-2 Receptor and Regulator of the Renin-Angiotensin System: Celebrating the 20th Anniversary of the Discovery of ACE2. Circ Res 126, 1456–1474.

Guo, M., Tao, W., Flavell, R.A., and Zhu, S. (2021). Potential intestinal infection and faecal-oral transmission of SARS-CoV-2. Nat Rev Gastroenterol Hepatol 18, 269–283.

Hadjadj, J., Yatim, N., Barnabei, L., Corneau, A., Boussier, J., Smith, N., Péré, H., Charbit, B., Bondet, V., Chenevier-Gobeaux, C., et al. (2020). Impaired type I interferon activity and inflammatory responses in severe COVID-19 patients. Science 369, 718–724.

Hao, Q., Dong, B.R., and Wu, T. (2015). Probiotics for preventing acute upper respiratory tract infections. Cochrane Database Syst Rev, Cd006895.

Harper, A., Vijayakumar, V., Ouwehand, A.C., ter Haar, J., Obis, D., Espadaler, J., Binda, S., Desiraju, S., and Day, R. (2021). Viral Infections, the Microbiome, and Probiotics. Frontiers in Cellular and Infection Microbiology 10.

Hashimoto, T., Perlot, T., Rehman, A., Trichereau, J., Ishiguro, H., Paolino, M., Sigl, V., Hanada, T., Hanada, R., Lipinski, S., et al. (2012). ACE2 links amino acid malnutrition to microbial ecology and intestinal inflammation. Nature 487, 477–481.

Hill, C., Guarner, F., Reid, G., Gibson, G.R., Merenstein, D.J., Pot, B., Morelli, L., Canani, R.B., Flint, H.J., Salminen, S., et al. (2014). Expert consensus document. The International Scientific Association for Probiotics and Prebiotics consensus statement on the scope and appropriate use of the term probiotic. Nat Rev Gastroenterol Hepatol 11, 506–514.

Hoagland, D.A., Moller, R., Uhl, S.A., Oishi, K., Frere, J., Golynker, I., Horiuchi, S., Panis, M., Blanco-Melo, D., Sachs, D., et al. (2021). Leveraging the antiviral type I interferon system as a first line of defense against SARS-CoV-2 pathogenicity. Immunity 54, 557–570 e555.

Hoffmann, M., Kleine-Weber, H., Schroeder, S., Kruger, N., Herrler, T., Erichsen, S., Schiergens, T.S., Herrler, G., Wu, N.H., Nitsche, A., et al. (2020). SARS-CoV-2 Cell Entry Depends on ACE2 and TMPRSS2 and Is Blocked by a Clinically Proven Protease Inhibitor. Cell 181, 271–280 e278.

Johnson, L.M., and Scott, P. (2007). STAT1 expression in dendritic cells, but not T cells, is required for immunity to Leishmania major. J Immunol 178, 7259–7266.

Kanehisa, M. (2019). Toward understanding the origin and evolution of cellular organisms. Protein Sci 28, 1947–1951.

Kanehisa, M., Furumichi, M., Sato, Y., Ishiguro-Watanabe, M., and Tanabe, M. (2021). KEGG: integrating viruses and cellular organisms. Nucleic Acids Res 49, D545–d551.

Kanehisa, M., and Goto, S. (2000). KEGG: kyoto encyclopedia of genes and genomes. Nucleic Acids Res 28, 27–30.

Kim, Y.I., Kim, S.G., Kim, S.M., Kim, E.H., Park, S.J., Yu, K.M., Chang, J.H., Kim, E.J., Lee, S., Casel, M.A.B., et al. (2020). Infection and Rapid Transmission of SARS-CoV-2 in Ferrets. Cell Host Microbe 27, 704–709 e702.

King, S., Glanville, J., Sanders, M.E., Fitzgerald, A., and Varley, D. (2014). Effectiveness of probiotics on the duration of illness in healthy children and adults who develop common acute respiratory infectious conditions: a systematic review and meta-analysis. Br J Nutr 112, 41–54.

Lee, J.G., Jaeger, K.E., Seki, Y., Wei Lim, Y., Cunha, C., Vuchkovska, A., Nelson, A.J., Nikolai, A., Kim, D., Nishimura, M., et al. (2021). Human CD36(hi) monocytes induce Foxp3(+) CD25(+) T cells with regulatory functions from CD4 and CD8 subsets. Immunology 163, 293–309.

Lee, S., Channappanavar, R., and Kanneganti, T.D. (2020). Coronaviruses: Innate Immunity, Inflammasome Activation, Inflammatory Cell Death, and Cytokines. Trends Immunol 41, 1083–1099.

Lehtoranta, L., Latvala, S., and Lehtinen, M.J. (2020). Role of Probiotics in Stimulating the Immune System in Viral Respiratory Tract Infections: A Narrative Review. Nutrients 12.

Lenth, R.V., Buerkner, P., Herve, M., Love, J., Riebl, H., and Singmann, H. (2021). emmeans: Estimated Marginal Means, aka Least-Squares Means. R package version 1.6.1.

Love, M.I., Huber, W., and Anders, S. (2014). Moderated estimation of fold change and dispersion for RNA-seq data with DESeq2. Genome Biology 15, 550.

Lu, Q., Liu, J., Zhao, S., Gomez Castro, M.F., Laurent-Rolle, M., Dong, J., Ran, X., Damani-Yokota, P., Tang, H., Karakousi, T., et al. (2021). SARS-CoV-2 exacerbates proinflammatory responses in myeloid cells through C-type lectin receptors and Tweety family member 2. Immunity 54, 1304–1319 e1309.

Miettinen, M., Pietila, T.E., Kekkonen, R.A., Kankainen, M., Latvala, S., Pirhonen, J., Osterlund, P., Korpela, R., and Julkunen, I. (2012). Nonpathogenic Lactobacillus rhamnosus activates the inflammasome and antiviral responses in human macrophages. Gut Microbes 3, 510–522.

Morovic, W., Roos, P., Zabel, B., Hidalgo-Cantabrana, C., Kiefer, A., Barrangou, R., and Müller, V. (2018). Transcriptional and Functional Analysis of Bifidobacterium animalis subsp. lactis Exposure to Tetracycline. Applied and Environmental Microbiology 84, e01999–01918.

Munoz-Fontela, C., Dowling, W.E., Funnell, S.G.P., Gsell, P.S., Riveros-Balta, A.X., Albrecht, R.A., Andersen, H., Baric, R.S., Carroll, M.W., Cavaleri, M., et al. (2020). Animal models for COVID-19. Nature 586, 509–515.

Munshi, I., Khandvilkar, A., Chavan, S., Sachdeva, G., Mahale, S., and Chaudhari, U. (2021). An overview of preclinical animal models for SARS-CoV-2 pathogenicity. Indian Journal of Medical Research 153, 17–25.

Netea, M.G., Giamarellos-Bourboulis, E.J., Dominguez-Andres, J., Curtis, N., van Crevel, R., van de Veerdonk, F.L., and Bonten, M. (2020). Trained Immunity: a Tool for Reducing Susceptibility to and the Severity of SARS-CoV-2 Infection. Cell 181, 969–977.

Patro, R., Duggal, G., Love, M.I., Irizarry, R.A., and Kingsford, C. (2017). Salmon provides fast and bias-aware quantification of transcript expression. Nature Methods 14, 417–419.

Peacock, T.P., Goldhill, D.H., Zhou, J., Baillon, L., Frise, R., Swann, O.C., Kugathasan, R., Penn, R., Brown, J.C., Sanchez-David, R.Y., et al. (2021). The furin cleavage site in the SARS-CoV-2 spike protein is required for transmission in ferrets. Nat Microbiol.

Ryan, K.A., Bewley, K.R., Fotheringham, S.A., Slack, G.S., Brown, P., Hall, Y., Wand, N.I., Marriott, A.C., Cavell, B.E., Tree, J.A., et al. (2021). Dose-dependent response to infection with SARS-CoV-2 in the ferret model and evidence of protective immunity. Nature Communications 12, 81.

Schultze, J.L., and Aschenbrenner, A.C. (2021). COVID-19 and the human innate immune system. Cell 184, 1671–1692.

Shi, H.Y., Zhu, X., Li, W.L., Mak, J.W.Y., Wong, S.H., Zhu, S.T., Guo, S.L., Chan, F.K.L., Zhang, S.T., and Ng, S.C. (2021). Modulation of gut microbiota protects against viral respiratory tract infections: a systematic review of animal and clinical studies. European Journal of Nutrition.

Stanifer, M.L., Kee, C., Cortese, M., Zumaran, C.M., Triana, S., Mukenhirn, M., Kraeusslich, H.G., Alexandrov, T., Bartenschlager, R., and Boulant, S. (2020). Critical Role of Type III Interferon in Controlling SARS-CoV-2 Infection in Human Intestinal Epithelial Cells. Cell Rep 32, 107863.

Taefehshokr, N., Taefehshokr, S., Hemmat, N., and Heit, B. (2020). Covid-19: Perspectives on Innate Immune Evasion. Front Immunol 11, 580641.

Turner, R.B., Woodfolk, J.A., Borish, L., Steinke, J.W., Patrie, J.T., Muehling, L.M., Lahtinen, S., and Lehtinen, M.J. (2017). Effect of probiotic on innate inflammatory response and viral shedding in experimental rhinovirus infection - a randomised controlled trial. Benef Microbes 8, 207–215.

Weiss, G., Rasmussen, S., Zeuthen, L.H., Nielsen, B.N., Jarmer, H., Jespersen, L., and Frokiaer, H. (2010). Lactobacillus acidophilus induces virus immune defence genes in murine dendritic cells by a Toll-like receptor-2-dependent mechanism. Immunology 131, 268–281.

Verdecchia, P., Cavallini, C., Spanevello, A., and Angeli, F. (2020). The pivotal link between ACE2 deficiency and SARS-CoV-2 infection. Eur J Intern Med 76, 14–20.

West, N.P., Horn, P.L., Pyne, D.B., Gebski, V.J., Lahtinen, S.J., Fricker, P.A., and Cripps, A.W. (2014). Probiotic supplementation for respiratory and gastrointestinal illness symptoms in healthy physically active individuals. Clin Nutr 33, 581–587.

Wickham, H. (2016). ggplot2: Elegant Graphics for Data Analysis. (New York: Springer-Verlag).

Vignesh, R., Swathirajan, C.R., Tun, Z.H., Rameshkumar, M.R., Solomon, S.S., and Balakrishnan, P. (2021). Could Perturbation of Gut Microbiota Possibly Exacerbate the Severity of COVID-19 via Cytokine Storm? Frontiers in Immunology 11.

Voichita, C., Ansari, S., and Draghici, S. (2019). ROntoTools: R Onto-Tools suite. In R package.

Yang, D., Chu, H., Hou, Y., Chai, Y., Shuai, H., Lee, A.C.-Y., Zhang, X., Wang, Y., Hu, B., Huang, X., et al. (2020). Attenuated Interferon and Proinflammatory Response in SARS-CoV-2–Infected Human Dendritic Cells Is Associated With Viral Antagonism of STAT1 Phosphorylation. The Journal of Infectious Diseases 222, 734–745.

Zabel, B.E., Gerdes, S., Evans, K.C., Nedveck, D., Singles, S.K., Volk, B., and Budinoff, C. (2020). Strain-specific strategies of 21.-fucosyllactose, 3-fucosyllactose, and difucosyllactose assimilation by Bifidobacterium longum subsp. infantis Bi-26 and ATCC 15697. Scientific Reports 10, 15919.

Zheng, J., Wang, Y., Li, K., Meyerholz, D.K., Allamargot, C., and Perlman, S. (2021). Severe Acute Respiratory Syndrome Coronavirus 2-Induced Immune Activation and Death of Monocyte-Derived Human Macrophages and Dendritic Cells. J Infect Dis 223, 785–795.

Zhou, R., To, K.K., Wong, Y.C., Liu, L., Zhou, B., Li, X., Huang, H., Mo, Y., Luk, T.Y., Lau, T.T., et al. (2020a). Acute SARS-CoV-2 Infection Impairs Dendritic Cell and T Cell Responses. Immunity 53, 864–877 e865.

Zhou, Z., Ren, L., Zhang, L., Zhong, J., Xiao, Y., Jia, Z., Guo, L., Yang, J., Wang, C., Jiang, S., et al. (2020b). Heightened Innate Immune Responses in the Respiratory Tract of COVID-19 Patients. Cell Host Microbe 27, 883–890 e882.

