## Supplemental Figures and Tables for "Probiotic consortia improve anti-viral immunity to SARS-CoV-2 in Ferrets"

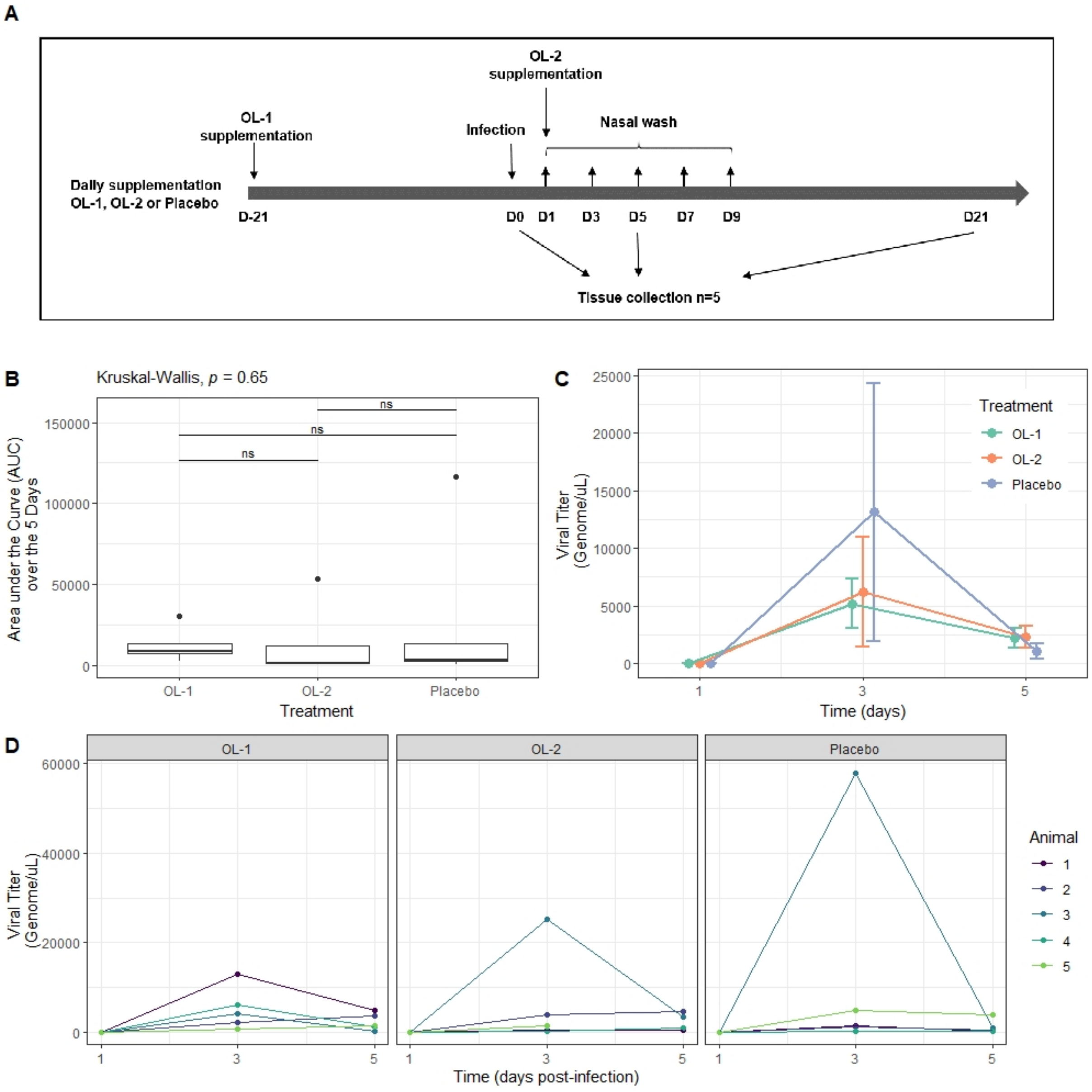


**Supplementary Figure 1. The viral titer analysis of the pilot ferret study.** The ferrets were supplemented with OL-1 or OL-2 or placebo and infected with SARS-CoV-2. (A) The ferret study design and time points for supplementation, SARS-CoV-2 infection, tissue and nasal wash collection; B) The nasal wash viral titer AUC analysis (Line: median Box: lower and upper quartile, Whiskers: min and max; Kruskal-Wallis test and pairwise Wilcoxon tests) C) The nasal wash viral titer timepoint analysis (Symbol: mean; error bars: standard error) D) Time course of individual viral titers in nasal washes from ferrets.

**Supplementary Table 1 Differential genes of significance**

Provided as separate excel-file.

**Supplementary Table 2 Pathway analysis**

Provided as separate excel-file.

**Supplementary Table 3 qPCR**

| Gene Target | Primer/probe | Sequences (5'-3') |
|---|---|---|
| ACE2 | Forward | GGAGCAAAGACGATGGATGT |
| | Reverse | TTTTGCTCTTTCAGCCAGGT |
| | Probe | CCACTGCTCAACTACTTCGAGCCCT |
| CCL2 | Forward | CCGGTCACCTGCTGCTATAC |
| | Reverse | ACTTGCTGCTGGTGACTCTC |
| | Probe | AGATCTCAGTGCAGAGGCTGGTGA |
| CCL5 | Forward | ATGCCAGCGGTCGTCTTTG |
| | Reverse | CGCACCCATTTCTTCTGTGG |
| | Probe | CACCGCCAAGTGTGTGCCAAC |
| CXCL10 | Forward | TCTGTCCGGGCACTATAAGC |
| | Reverse | GAGGAGCGTGTCAGTAGCAG |
| | Probe | TAGTGAACGTACCAGGTCCTAGCC |
| IL1β | Forward | TGAAGCTGCAGTTCTGACGA |
| | Reverse | AAAGGCGTGGAGTGGGTTTT |
| | Probe | AGTTCAGAACTCCGATGCCAGTGC |
| IL6 | Forward | CAAGTGGCTGAAACACGTAACAA |
| | Reverse | GGCTGAACTGCAGGAAATCC |
| | Probe | TCACCTCATCCTACGGAGCCTTG |
| IL8 | Forward | TGAAGCTGCAGTTCTGACGA |
| | Reverse | AAAGGCGTGGAGTGGGTTTT |
| | Probe | AGTTCAGAACTCCGATGCCAGTGC |
| IL10 | Forward | CGAGAACCACGACCCAGAA |
| | Reverse | CCGCAGGGTCTTCAGCTTT |
| | Probe | TCAAGGAGCACGTGAACTCGCTGG |
| IFNα | Forward | AGACCTTCCACCTCTTCTGC |
| | Reverse | GTCCTGAGCACAATTCCTCCA |
| | Probe | CTCACCTGCTCCCTGGAACACGAC |
| IFNγ | Forward | TGGTGGGCCTCTTTTCTTAGATAT |
| | Reverse | AGAAGGAGACAATTTGGCTTTGA |

| | | |
|---|---|---|
| | Probe | TTGAAGAACTGGAGAGAGGAGAGTGACAAAAAAA |
| IFNL1 | Forward | GCTGTGACATTGGCAGGTTC |
| | Reverse | CCAAAGCATCCTTGGCCATC |
| | Probe | AATCTCTGTCACCACGGGAGCTG |
| TLR8 | Forward | ACAAAATGGCTCTGTGATTGCA |
| | Reverse | TGTCACATACTTGCCCACGG |
| | Probe | CCGTCGACTGAGAGAAGTTCCCCA |
| TNFα | Forward | ATGTTGTAGCAAACCCTGAAGCT |
| | Reverse | ATTGGCCAGGAGGGCATT |
| | Probe | ACTCCAATGGCTGAGCCGACGTG |
| RPS18 | Forward | TCACCAAGAGAGCCGGAGAG |
| | Reverse | TTGGCGAGGGTTCTGCATAA |
| | Probe | ACTGAGGATGAGGTGGAACGTGT |
| IFIH1 | Forward | CTGGGACGAGGACAATGACT |
| | Reverse | AGCAGCTGACACTTCCTTCT |
| | Probe | ACCGGACTCGTCTTTGGTGGATTCT |
| NFkB | Forward | CCTTCCAGTTTCTCTCCATTTGT |
| | Reverse | TCCTTTTCATTATCTTGGGCCT |
| | Probe | AGAAAACGCAGCAGTGGAAATTGC |
| GAPDH | Forward | CTGCTGATGCCCCCATGT |
| | Reverse | TTGCTGACAATCTTGAGGGAGTT |
| | Probe | TCATACTTCTCATGGTTCACACCCATCACG |

**Supplementary Table 4 Transcriptomics QC data**

| sample | Cell type | condition | Donor | Num processed (millions) | Num mapped (millions) | Percent mapped |
|---|---|---|---|---|---|---|
| 1 | DC | OL-1 | D4 | 7.2 | 5.9 | 82.1 |
| 2 | DC | OL-1 | D1 | 4.5 | 3.9 | 87.2 |
| 3 | DC | OL-1 | D2 | 6.0 | 4.4 | 72.6 |
| 4 | DC | OL-1 | D3 | 7.1 | 6.3 | 89.6 |
| 5 | DC | OL-2 | D4 | 7.7 | 6.6 | 86.2 |
| 6 | DC | OL-2 | D1 | 4.3 | 3.5 | 80.7 |
| 7 | DC | OL-2 | D2 | 5.9 | 4.1 | 69.2 |
| 8 | DC | OL-2 | D3 | 7.6 | 6.6 | 86.6 |
| 9 | DC | ctrl | D4 | 7.7 | 5.8 | 75.1 |
| 10 | DC | ctrl | D1 | 8.1 | 5.9 | 72.6 |
| 11 | DC | ctrl | D2 | 7.4 | 4.5 | 61.4 |
| 12 | DC | ctrl | D3 | 8.4 | 7.0 | 83.2 |
| 13 | DC | polyIC+R848 | D4 | 9.8 | 8.7 | 88.0 |
| 14 | DC | polyIC+R848 | D1 | 5.3 | 3.6 | 67.2 |
| 15 | DC | polyIC+R848 | D2 | 8.4 | 6.2 | 73.4 |
| 16 | DC | polyIC+R848 | D3 | 8.3 | 7.1 | 85.4 |
| 17 | Mf | OL-1 | D4 | 12.3 | 10.5 | 85.5 |
| 18 | Mf | OL-1 | D1 | 9.7 | 8.6 | 88.1 |
| 19 | Mf | OL-1 | D2 | 9.9 | 8.6 | 86.8 |
| 20 | Mf | OL-1 | D3 | 10.9 | 8.9 | 81.8 |
| 21 | Mf | OL-2 | D4 | 7.4 | 6.3 | 85.0 |
| 22 | Mf | OL-2 | D1 | 5.8 | 4.8 | 81.8 |
| 23 | Mf | OL-2 | D2 | 7.4 | 6.3 | 84.0 |
| 24 | Mf | OL-2 | D3 | 5.9 | 5.0 | 83.7 |
| 25 | Mf | ctrl | D4 | 8.2 | 6.3 | 76.7 |
| 26 | Mf | ctrl | D1 | 9.9 | 8.6 | 86.2 |
| 27 | Mf | ctrl | D2 | 8.1 | 7.0 | 86.5 |
| 28 | Mf | ctrl | D3 | 9.9 | 8.2 | 82.4 |
| 29 | Mf | polyIC+R848 | D4 | 8.4 | 7.2 | 85.9 |
| 30 | Mf | polyIC+R848 | D2 | 8.3 | 7.1 | 84.9 |
